# Differences in directed functional brain connectivity related to age, sex and mental health

**DOI:** 10.1101/799593

**Authors:** Martina J. Lund, Dag Alnæs, Simon Schwab, Dennis van der Meer, Ole A. Andreassen, Lars T. Westlye, Tobias Kaufmann

## Abstract

**Objective:** Functional interconnections between brain regions define the ‘connectome’ which is of central interest for understanding human brain function. Resting-state functional magnetic resonance (rsfMRI) work has revealed changes in static connectivity related to age, sex, cognitive abilities and psychiatric symptoms, yet little is known how these factors may alter the information flow. The commonly used approach infers functional brain connectivity using stationary coefficients yielding static estimates of the undirected connection strength between brain regions. Dynamic graphical models (DGMs) are a multivariate model with dynamic coefficients reflecting directed temporal associations between nodes, and can yield novel insight into directed functional connectivity. Here, we leveraged this approach to test for associations between edge-wise estimates of direction flow across the brain connectome and age, sex, intellectual abilities and mental health.

**Methods:** We applied DGM to investigate patterns of information flow in data from 984 individuals from the Human Connectome Project (HCP) and 10,249 individuals from the UK Biobank.

**Results:** Our analysis yielded patterns of directed connectivity in independent HCP and UK Biobank data similar to those previously reported, including that the cerebellum consistently receives information from other networks. We show robust associations between information flow and age and sex for several connections, with strongest effects of age observed in the sensorimotor network. Visual, auditory and sensorimotor nodes were also linked to mental health.

**Discussion:** Our findings support the use of DGM as a measure of directed connectivity in rsfMRI data and provide new insight into the shaping of the connectome during aging.

## Introduction

Although the rates and trajectories vary substantially between individuals and cognitive domains (Ardila, 2007), normal aging is primarily associated with a decline in most cognitive functions, including executive functions, attention, memory and perception (Riddle, 2007). Numerous studies have established pronounced age-related differences in brain network connections (Betzel et al., 2014; Cassady et al., 2019; Dørum et al., 2017; Geerligs, Renken, Saliasi, Maurits, & Lorist, 2015; Grady, Springer, Hongwanishkul, McIntosh, & Winocur, 2006; Maglanoc, Kaufmann, van der Meer, et al., 2019; Meunier, Achard, Morcom, & Bullmore, 2009; Wang, Su, Shen, & Hu, 2012). However, so far mostly age-related network changes have been studied using static functional connectivity, where undirected connectivity strengths are estimated from stationary coefficients and assumed not to change short-term during the period of scan. Dynamic functional connectivity (i.e., time-varying connectivity) has been studied to a lesser degree yet could yield new knowledge about connectivity direction, thereby supplementing approaches for static connectivity with insight into the information flow of neural activity, underlying processes related to cognitive functions and mental health (Hutchison et al., 2013).

There are various approaches to estimate connectivity direction, often divided into *effective connectivity* and *directed functional connectivity* (K. Friston, Moran, & Seth, 2013). Effective connectivity refers to the causal influence that one node exerts over another (Bielczyk et al., 2019; K. J. Friston, 2011), while directed functional connectivity (dFC) denotes information flow between nodes by estimating statistical interdependence using measured blood-oxygen-level-dependent (BOLD) responses (Bielczyk et al., 2019). Recent work has provided evidence of changes in connectivity direction with age. For instance, one study noted posture-related changes in effective connectivity with elderly, compared to younger participants showing higher effective connectivity between the prefrontal cortex (PFC) and the motor cortex (MC) as measured using functional near-infrared spectroscopy (fNIRS) while standing (Huo et al., 2018). Studies have also reported age-related psychomotor slowing with higher effective connectivity (Michely et al., 2018), in addition to changes in effective connectivity during resting-state functional magnetic resonance imaging (rsfMRI) in certain areas of the brain of elderly APOE ε4 carriers (Luo et al., 2019). It has also been shown that there are alterations in effective connectivity in the prefrontal cortex during emotion processing in individuals with autism spectrum disorders (Wicker et al., 2008), and disrupted effective connectivity in patients with externalizing behavior disorders (Shannon, Sauder, Beauchaine, & Gatzke-Kopp, 2009), schizophrenia (Schlösser et al., 2003) and depression (Lu et al., 2012; Rolls et al., 2018). Others have investigated effective connectivity in rsfMRI in relation to psychedelics and found evidence for alterations in cortico-striato-thalamic-cortico loops in individuals given LSD (Preller et al., 2019). Changes in effective connectivity have also been observed in relation to episodic simulation and social cognition (Pehrs, Zaki, Taruffi, Kuchinke, & Koelsch, 2018), as well as memory function in a neurodevelopmental sample (Riley et al., 2018). However, we know little about how the flow of information between different brain networks is altered throughout life and how this affects the vulnerability for mental disorders. Gaining such knowledge is of importance given the known associations between age-related brain changes and mental disorders (Kaufmann et al., 2019; Koutsouleris et al., 2013; Schnack et al., 2016). In addition, it is unknown how sex differences contribute to differences in information flow. New statistical methods such as Dynamic graphical models (DGM) allow us to examine the extent to which developmental and age-related processes in the brain change the connection between networks, and how this is associated with various factors such as sex, cognitive abilities and mental health.

DGM has been proposed as an approach for estimating dFC in rsfMRI data and explore intrinsic functional connectivity in relation to organization of brain functioning. DGM is a form of Dynamic Bayesian Networks, that describes the instantaneous directed relationships between nodes (Bilmes, 2010; Schwab et al., 2018). As such, it does not make assumptions based solely on lagged relationships which can be highly influenced by the hemodynamic response, such as for instance Granger Causality assumes. Further, DGM utilizes dynamic linear models (DLMs) for each node or network to estimate binary relationships. While DLMs are acyclic, the overall network can be cyclic. From this, one can study the spatiotemporal arrangement of links in the network, defined here as the directionality between a node pair. Accordingly, this statistical method can give a meaningful characterization of the dynamic connectivity between network nodes.

Initial implementation of the DGM approach in rsfMRI data from mice (N=16) showed information flow from CA1 and dentate gyrus to the cingulate cortex, which is in line with studies that have used viral tracers to examine the directed anatomical connectivity (Schwab et al., 2018). Further, human rsfMRI data in Human Connectome Project (HCP) subjects (N=500) suggested consistent default mode network (DMN) influence on cerebellar, limbic and auditory/temporal networks, in addition to a stable mutual relationship between visual medial (VM) and visual lateral (VL) networks (Schwab et al., 2018). Accordingly, DGM is a promising method to further disentangle the functional connectome. Here, we aimed to replicate findings from Schwab et al. (2018) using independent data from the HCP (Van Essen et al., 2013) and the UK Biobank (Sudlow et al., 2015). Further, we aimed to investigate if there were associations between dFC and age, sex, intellectual abilities and mental health measures, assessing mental health as a continuum in *healthy* (undiagnosed) individuals. We tested these associations for every connection of the directed network (edge-level analysis), and on node-level by assessing associations with network in-degree (the number of input connections for a given node) and out-degree (the number of output connections for a given node). Overall, we expected to find alterations in information flow to prefrontal areas with higher age as studies have shown large scale reorganization of the brain with pronounced effects in frontal regions (Huo et al., 2018; Luo et al., 2019; Michely et al., 2018).

## Methods

### Study samples

HCP: The HCP consortium is funded by the National Institutes of Health (NIH) led by Washington University, University of Minnesota, and Oxford University. HCP is undertaking a systematic effort to map macroscopic human brain circuits and their relationship to behavior in a large population of young healthy adults (Van Essen et al., 2013). HCP participants are drawn from a healthy population born in Missouri, where a proportion of the subjects included are adult twins and their non-twin siblings (Van Essen et al., 2013). The adult sample consists of 1200 subjects. Exclusion criteria include having siblings with severe neurodevelopmental disorders, and documented neuropsychiatric or neurologic disorders. Furthermore, individuals with illnesses such as diabetes or high blood pressure and twins born prior to 34 weeks’ gestation and non-twins born prior to 37 weeks’ gestation were excluded (Van Essen et al., 2013). The participants went through an MRI protocol, in addition to extensive behavioral assessment outside the scanner, in the domains of cognitive, emotional, motor, and sensory functions (Van Essen et al., 2013). All participants provided signed informed consent. Washington University Institutional Review Board approved the study (Glasser et al., 2016).

UK Biobank: The UK Biobank initiative is a large-scale biobank prospective cohort established by the Medical Research Council and Wellcome Trust (Collins, 2012), and funded by the UK Medical Research Council, Wellcome Trust, Department of Health, British Heart Foundation, Diabetes UK, Northwest Regional Development Agency, Scottish Government, and Welsh Assembly Government (Sudlow et al., 2015). This population-based study examines the influence of genetic and environmental factors and the occurrence of disease in participants included in the age range of 40-69 years old, recruited from 2006-2010, and were 45-80 years when they were scanned in the years thereafter (Sudlow et al., 2015). The study has recruited 500 000 subjects, where 100 000 are going to be included as an MRI subgroup (Miller et al., 2016). Further, participants filled out questionnaires about lifestyle, family, as well as medical history in addition to completing a variety of physical measures (Sudlow et al., 2015). In addition, a subset of participants filled in a mental health questionnaire (MHQ) online. All participants provided signed informed consent. UK Biobank was approved by the National Health Service National Research Ethics Service (ref 11/NW/0382, (Health Research Authority, 2016)).

### MRI acquisition

MR data was collected by the study teams of HCP and UK Biobank.

#### HCP

MRI data from the HCP study was collected using a customized 3T Siemens Skyra with a 32-channel receive head coil at Washington University, US. Resting-state blood-oxygen-level-dependent (BOLD) fMRI data was collected for each subject using a T2*-weighted BOLD echo-planar imaging (EPI) sequence with the following parameters: repetition time (TR)/echo time (TE)/flip angle (FA) = 720ms/33.1ms/52°; voxel size, 2.0×2.0×2.0 mm, MB=8, BW = 2290 Hz/Px, in-plane FOV = 208 × 180 mm, fat sat, 1200 volumes; scan time ≈ 15min (Smith et al., 2013). A T1-weighted 3D MPRAGE, sagittal sequence with the following pulse sequence parameters was obtained: TR/TE/FA = 2.4ms/2.14ms/8**°**; voxel size = 0.7 × 0.7 × 0.7 mm, FOV: 88×224×224, iPAT=2, scan time = 7min 40 sec. The T1-weighted image was used for registration to the EPI data in the present study. rsfMRI data were collected over 2 days divided into 4 rsfMRI sessions where the scanning session took 1 hour each of the days, including task fMRI (Glasser et al., 2016).

#### UK Biobank

MR data from the UK Biobank study was collected with a 3T standard Siemens Skyra using a 32-channel receive head coil at Newcastle and Cheadle Imaging Centre in the UK. Resting-state fMRI data was collected for each subject using a T2*-weighted BOLD echo-planar imaging (EPI) sequence with the following parameters: TR/TE/FA = 735ms/39ms/52°; voxel size, 2.4×2.4×2.4 mm, MB=8, R=1, no iPAT, fat sat, 490 volumes; scan time = 6min 10 sec. A T1-weighted 3D MPRAGE, sagittal sequence with the following pulse sequence parameters was obtained: TR/TE/FA = 2.0ms/2.01ms/8**°**; voxel size = 1.0 × 1.0 × 1.0 mm, FOV: 208×256×256, in-plane acceleration iPAT=2, scan time = 5 min. The T1-weighted image was used for registration to the EPI data in the present study. The entire MRI protocol took 31 minutes in effective scan time (Miller et al., 2016).

### MRI preprocessing

#### HCP

Processed HCP data was obtained from the HCP database (https://ida.loni.usc.edu/login.jsp), where we downloaded the released PTN 1200-subjects package. The HCP project processed the data through their pipeline, which is specifically made for HCP high-quality data (Glasser et al., 2013). Their preprocessing comprised image processing tools, based on Smith et al. (2013), with minimal-preprocessing according to Glasser et al. (2013). In addition, areal-feature-based alignment and the multimodal surface matching algorithm was applied for inter-subject registration of the cerebral cortex (Glasser et al., 2013; Robinson et al., 2014). Further, artefacts were removed by means of FIX (FMRIB’s ICA-based X-noisiefier, (Griffanti et al., 2014; Salimi-Khorshidi et al., 2014)), and ICA (independent component analysis, (Beckmann & Smith, 2004)) while dual regression was used for further processing of timeseries, these steps are described in more detail below. HCP structural data was manually quality checked while the fMRI data went through a built in quality control pipeline where estimates including voxel-wise temporal standard deviation (tSD), temporal SNR (tSNR), movement rotation and translation were computed (Marcus et al., 2013). In addition, the BIRN Human QA tool was used (Glover et al., 2012; Marcus et al., 2013). 184 subjects were reconstructed using an earlier version of the HCP data reconstruction software, while 812 subjects were run through a later edition, and 7 subjects was processed using a mixture of the two methods. Further, the data was temporally demeaned and variance normalized (Beckmann & Smith, 2004). Next, fMRI datasets were submitted to a group ICA, a data driven analysis technique used to discover independently distributed spatial patterns that represent source processes in the data (Beckmann & Smith, 2004). ICA extracts spatially independent components, a set of spatial maps and associated time courses, by use of blind signal source separation and linear decomposition of fMRI data (McKeown et al., 1998; McKeown & Sejnowski, 1998). MIGP (MELODIC’s Incremental Group-PCA) from 468 subjects were used to generate group-PCA that was used for the group-ICA utilizing FSL’s Multivariate Exploratory Linear Optimized Decomposition into Independent Components (MELODIC) tool (Beckmann & Smith, 2004; Hyvärinen, 1999), where 25 components were extracted and used for further processing. ICA was applied in grayordinate space (Glasser et al., 2013). Dual regression was applied to estimate specific spatial maps and corresponding time series from the group ICA for each subject (Beckmann & Smith, 2004; Filippini et al., 2009). As Schwab et al. (2018) reported a high degree of consistency in dFC patterns between rsfMRI sessions, we included data from the first run in our analysis and as follows this was used for further processing where dual regression was applied.

#### UK Biobank

Processed data was accessed from the UK Biobank study team under accession code 27412. The Biobank preprocessing comprised image processing tools, largely acquired from FSL (http://fsl.fmrib.ox.ac.uk), and complied with the pre-processing steps done as part of the HCP pipeline, including motion correction using MCFLIRT, grand-mean intensity normalisation of the 4D dataset by a single multiplicative factor, high pass temporal filtering and distortion correction (Alfaro-Almagro et al., 2018). The EPI unwarping step included alignment to the T1, where the unwarped data is written out in native fMRI space, while the transform to T1 space is written out independently (Alfaro-Almagro et al., 2018). Fieldmaps were utilized as part of the melodic pipeline (Alfaro-Almagro et al., 2018). FMRIB’s Linear Image Registration tool (FLIRT) was used to register fMRI volumes to the T1-weighted image (Mark Jenkinson, Bannister, Brady, & Smith, 2002; M. Jenkinson & Smith, 2001). Boundary based registration (Greve & Fischl, 2009) was used in a final step to refine the registration of the EPI and structural image. The ICA+FIX and dual regression procedure corresponds to what we reported for HCP above. For the UK Biobank sample, 4100 fMRI datasets were submitted to a group ICA, where 25 components where extracted from the ICA and used for further analysis. A FIX classifier for UK Biobank imaging data was hand trained on 40 Biobank rsfMRI datasets for removal of artefacts (Alfaro-Almagro et al., 2018). As for quality assessment, part of the UK Biobank imaging pipeline entails assessment of the T1-weighted images, which includes automated classification by use of machine learning (Alfaro-Almagro et al., 2018). If a T1-weighted image has been classified as having serious issues, the dataset has not been used in this study.

### Included participant data

#### HCP

From the HCP data release, four subjects were excluded due to missing information about mean relative motion and 15 individuals were excluded due to missing information on cognitive or mental health data, yielding data from a total of 984 individuals aged 22-37 years (mean: 28.7 years, sd: 3.71 years, 52.8% females) for the analysis on all HCP subjects. Out of those, data from 495 individuals were not included by Schwab et al. (2018) and were included for an additional replication analysis (mean: 28.6 years, sd: 3.72 years, 49.5% females).

#### UK Biobank

From the UK Biobank data release, we started out with 16,975 subjects, where we excluded subjects with a diagnosed neurological or psychiatric disorder (N=1,319) as well as 5,082 subjects missing information on mean relative motion, cognitive and mental health data, and 325 subjects that had a different number of volumes than in the standard protocol, yielding data from a total of 10,249 individuals aged 45-80 years (mean: 62.8 years, sd: 7.35 years, 53.8% females).

### Network analysis

Based on our aim to replicate findings from Schwab et al. (2018) in independent data, we chose the same model order as this study (d=25) for both HCP and the UKB sample. In each sample, we chose ten resting-state networks (RSNs) that had the highest spatial correlation with the ten RSNs reported by Smith et al. (2009), and in line with the procedure used in Schwab et al. (2018). These RSNs comprised default mode (DMN), cerebellar (Cer), visual occipital (VO), visual medial (VM), visual lateral (VL), right frontoparietal (FPR), left frontoparietal (FPL), sensorimotor (SM), auditory (Au), and executive control (Ex) networks. The timeseries for the ten RSNs were mean centered so that each timeseries for each node had a mean of zero. Finally, utilizing the DGM package v1.7.2 in R we estimated dFC from individual level RSN time series. RSNs will henceforth be referred to as network “nodes” as we estimated temporal connectivity between RSNs.

DGM is a graphical model with directed relationships between nodes and time-varying connectivity strengths for resting-state fMRI data and is a continuation of the Multiregression Dynamic Model (Costa et al., 2015; Queen & Smith, 1993). DGM comprises a set of dynamic linear models (DLMs), state space models that are linear and Gaussian (West and Harris, 1997). A single DLM is a directed acyclic graph (DAG), however, the DGM as a set of DLMs are allowed to contain cycles. In a DLM, the time series of a specified receiving node is regressed on the time series from one or more other transmitter nodes by deploying dynamic regression, where the directed relationship corresponds to information flow from the transmitter node to the receiver node (Schwab et al., 2018). Key steps of DGM include applying random walk smoothness for modelling the underlying coupling between time series, and deploying a Bayesian framework where dynamic directed graphical models (which includes time-varying coefficients for a set of transmitter nodes as covariates on a receiver node (Schwab et al., 2018)), gives a binary view of coupling, where a discount factor (δ) is given for each model. The “winning” model is selected based on the model evidence (derived from log-likelihood of the observed time-series), indicating if there is an influence or not from one transmitter node to a receiving node, and this resembles the approach used in regression dynamic causal modelling (Frässle et al., 2017).

### Statistical analysis

For both HCP and UK Biobank data, we performed logistic regression for every connection of the directed network using directed connectivity as the response variable and testing for associations with age, age-orthogonalized age squared (age^2^, using the poly function in R), sex, intellectual abilities, mental health, and motion taken together in one model for each sample. In the case of UK Biobank where data was acquired at multiple scanners, this model also included scanning site as a covariate. Figure 3 and 5a show the coefficients for the covariates extracted from the HCP - and the UK Biobank model per edge. We refer to this as edge-level analysis. Furthermore, we assessed input and output connections for a given network separately, to examine which nodes in general send and receive information. Accordingly, we calculated the number of output connections (denoted as out-degree) and the number of input connections (denoted as in-degree) for a given node, and we refer to this as node-level analysis. We performed linear regression using this in-degree and out-degree as dependent variables and the same independent variables as used on the edge-level. All p-values were Bonferroni corrected for a number of 90 analyses on the edge-level and for 10 analysis on the node-level, with an alpha level of 0.05.

**Figure 1:**
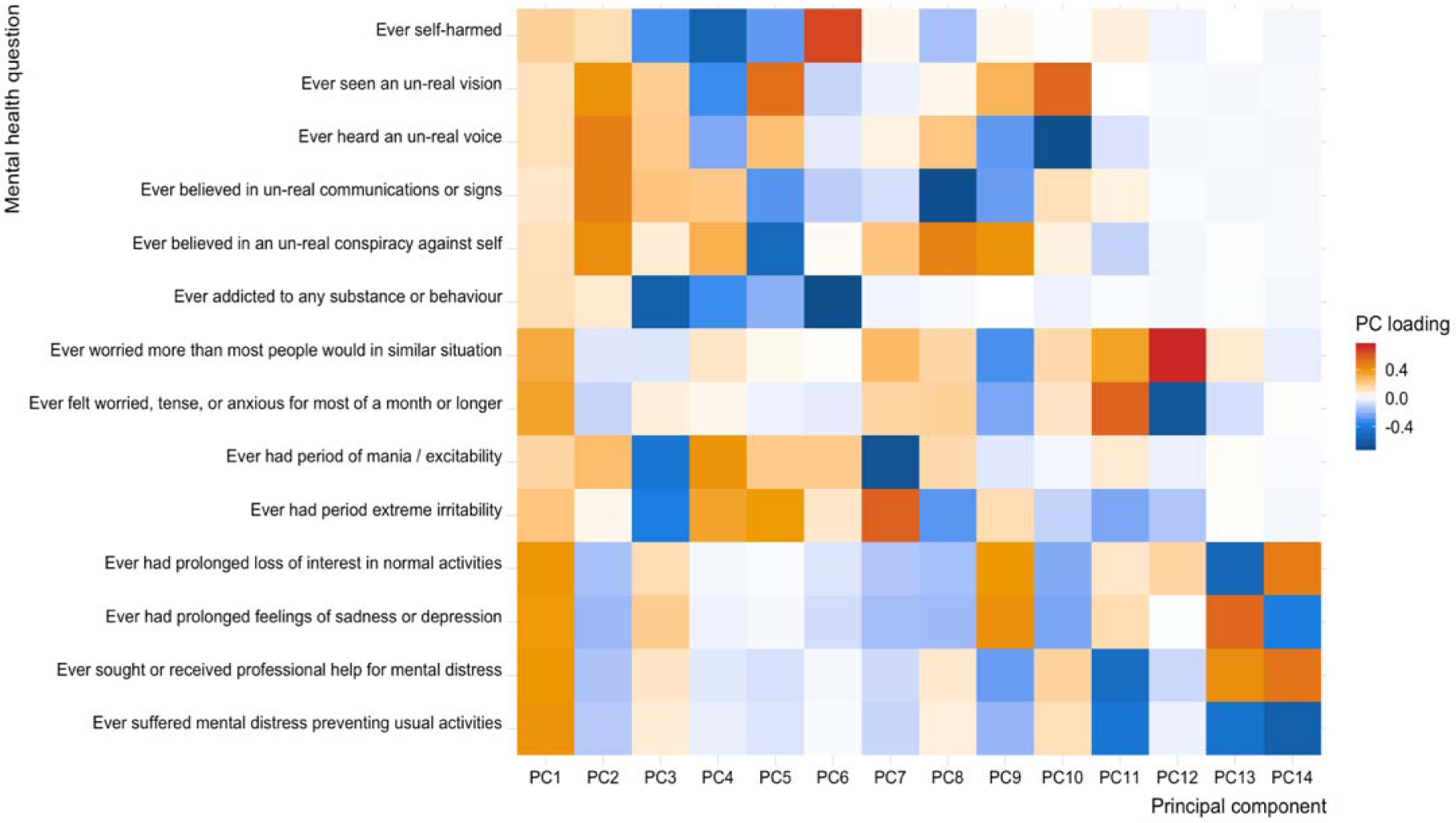
Principal component analysis (PCA) of mental health questionnaire from UK Biobank. We used the first two principal components as proxies of general psychopathology, referred to as “pF” and “pF_2_”.

**Figure 2:**
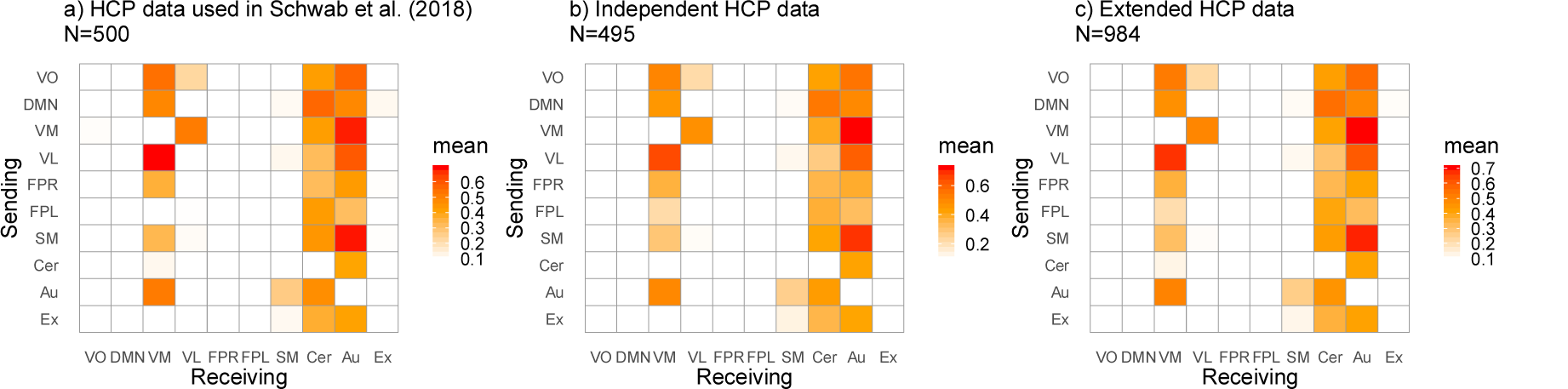
Average directed connectivity matrices across subjects for HCP data showing the significant proportions of edges (binomial test, 5% FDR threshold) in a) data previously reported by Schwab et al. (2018), b) independent data, c) all available data (a+b; slight differences in sample size due to differences in exclusion criteria). The legend shows the 10 RSNs included in the analysis; VO, visual occipital; DMN, default mode; VM, visual medial; VL, visual lateral; FPR, frontoparietal right; FPL, frontoparietal left; SM, sensorimotor; Cer, cerebellum; Au, auditory; Ex, executive control network, where the y-axis indicates the sender node, while the x-axis refers to the same nodes but here they are receivers.

**Figure 3:**
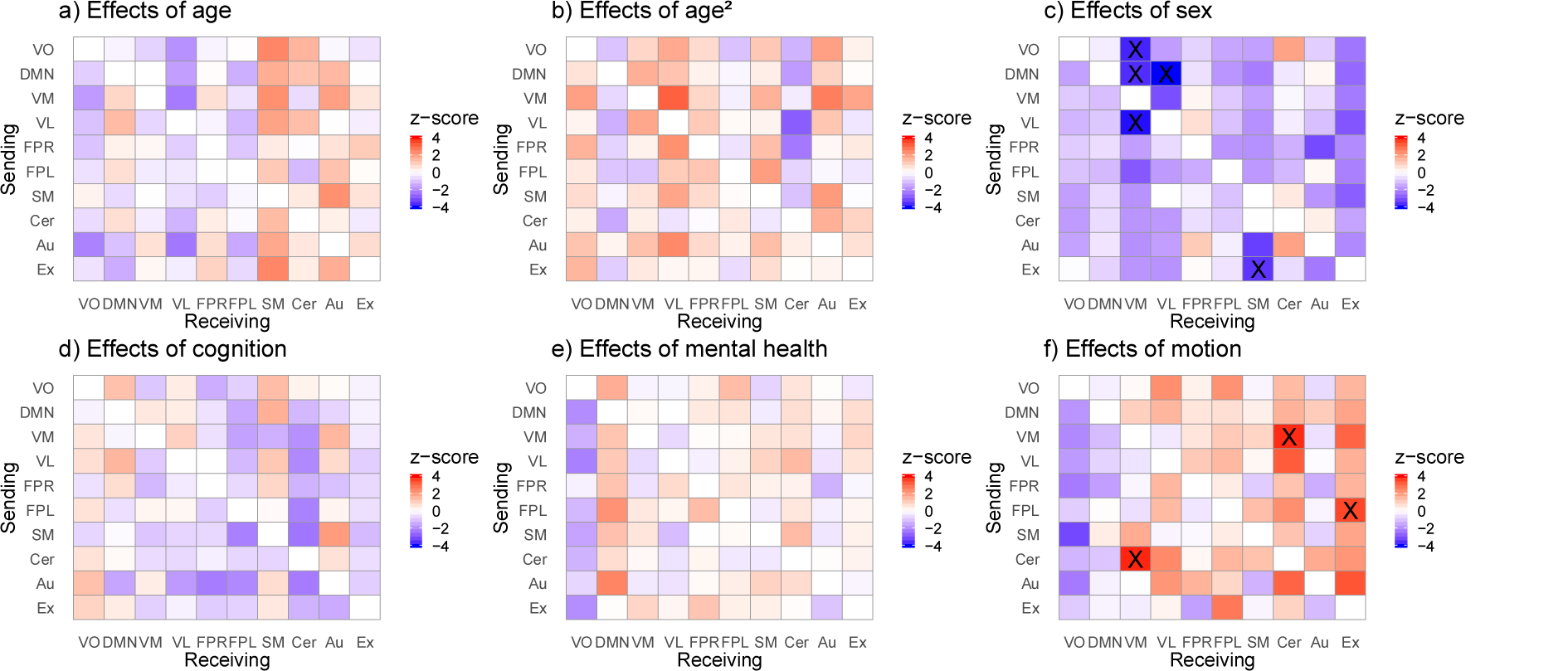
Directed connectivity matrices showing the effects of age (a), age^2^ (b), sex (c), intellectual abilities (d), mental health (e) and motion (f) on directed connectivity. The analysis was performed in all available HCP data (N=984, 22-37 years, df=977). Significant edges following Bonferroni correction are marked as X. The legend shows the 10 RSNs included in the analysis; VO, visual occipital; DMN, default mode; VM, visual medial; VL, visual lateral; FPR, frontoparietal right; FPL, frontoparietal left; SM, sensorimotor; Cer, cerebellum; Au, auditory; Ex, executive control network, where the y-axis indicates the sender node, while the x-axis refers to the receiving node. The colors reflect the z-value for the corresponding effects where red indicates a positive association and blue a negative association.

**Figure 4:**
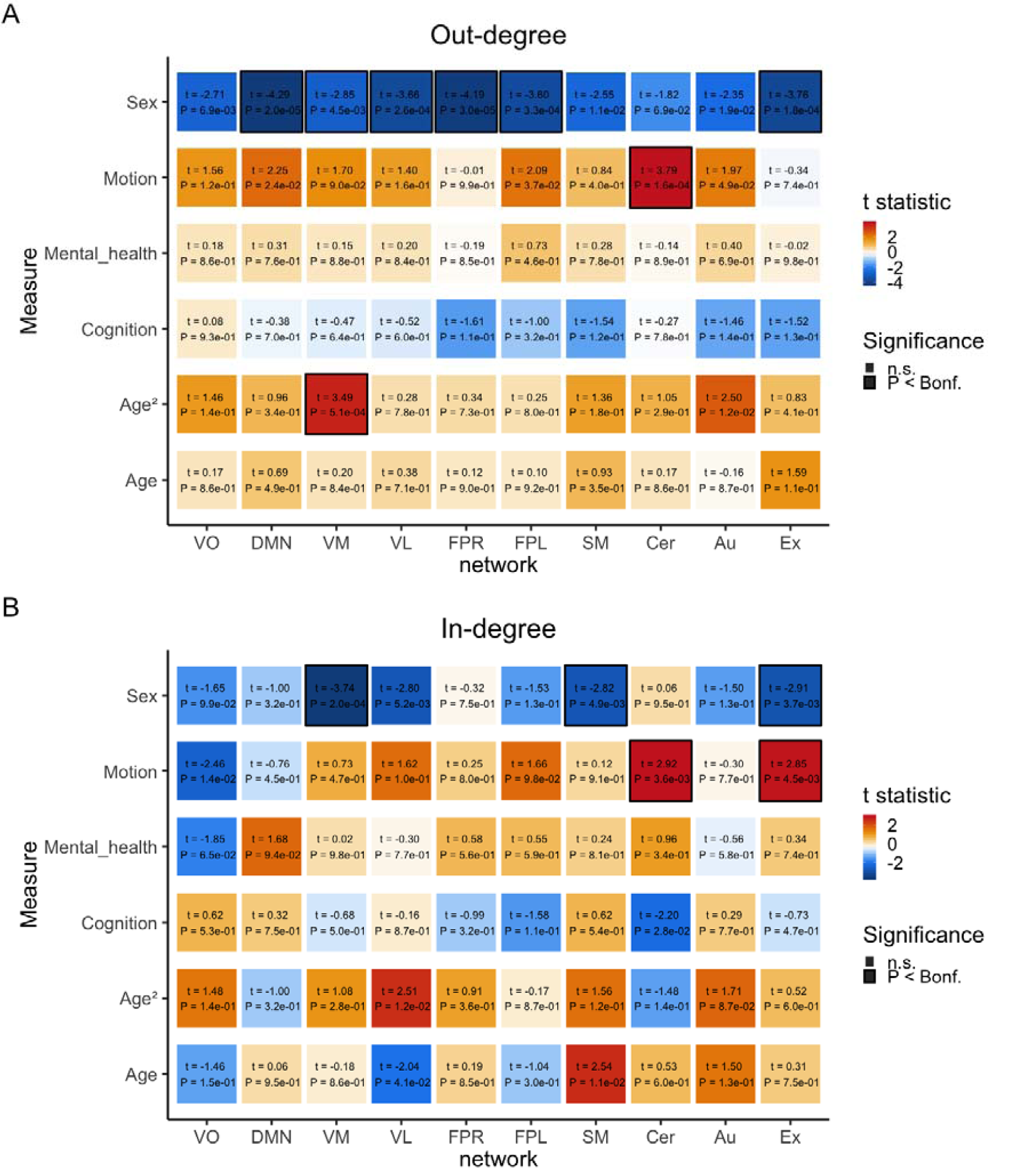
A) Out-degree matrix with corresponding effects of covariates age, age^2^, sex, cognition, mental health, and motion in HCP data (N=984, 22-37 years, df=977). B) In-degree matrix with corresponding effects for the same covariates as in panel A. The colors reflect the t-value for the corresponding effect where red indicates a positive association and blue a negative association. Numbers inside the boxes indicate t-statistic and p-value, where significant effects are marked with a black border following Bonferroni correction (p<0.05).

**Figure 5:**
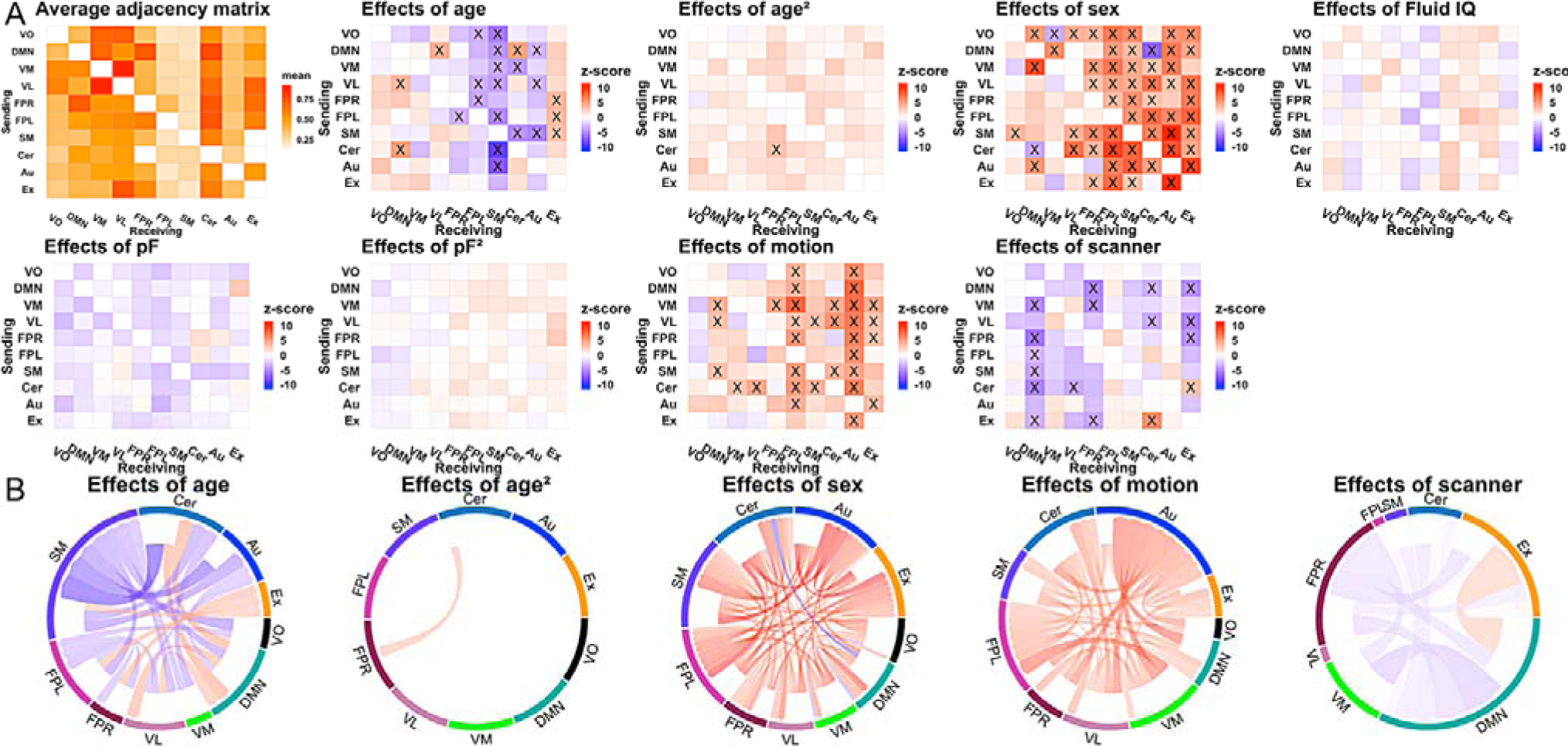
**A)** Average directed connectivity matrix showing significant proportion of edges (binomial test, 5% FDR threshold), and corresponding effects of age, age^2^, sex, fluid intelligence, pF, pF_2,_ motion and scanner for UK Biobank (N=10,249, 45-80 years, df=10240). Significant edges following Bonferroni correction are marked as X. **B)** chord diagrams that display only the significant effects of age, age^2^, sex, motion and scanner for the UK Biobank sample. The colors reflect the z-value for the corresponding effects where red indicates a positive association and blue a negative association. The arrow heads in the circular plots indicate direction (receiver or sender).

For the HCP data, we utilized the age-adjusted NIH Toolbox Cognition Total Composite Score as a measure of cognitive abilities which includes test in the following subdomains: executive function, episodic memory, language, processing speed and working memory (Barch et al., 2013). The gender and age adjusted T-score of the Achenbach Adult Self-Report, Syndrome Scales and DSM-Oriented Scale (ASR) was used as a measure of mental health for the HCP participants. In addition, we included the mean relative motion and the statistical models tested in HCP thus included age, age^2^, sex, cognitive test performance, mental health and motion.

For UK Biobank, we used the fluid intelligence score (UKB field: 20016, which consisted of the sum of the number of correct answers given to 13 fluid intelligence items) where we controlled for age on fluid intelligence before using the residuals in the analysis as a measure of cognitive abilities for participants in the UK Biobank sample. Further, we inferred mental health by performing a principal component analysis (PCA) on 14 items of the online MHQ available for 154,607 participants with less than 3 missing values on the included items (Figure 1). We imputed missing values in R using the missMDA package (Josse & Husson, 2016) and subsequently performed the PCA using the “prcomp” function. The first PC, often referred to as the p-Factor or pF (Caspi et al., 2013), explained 27.02% of the variance. This component related mostly to depression/anxiety items. Given recent indications that psychopathology may not be explained by a single dimension (Mallard et al., 2019), we also included the second principal component, which explained 11.94% of the variance. We refer to this component as pF_2,_ and this component related mostly to psychosis items. The statistical models tested in UK Biobank thus included age, age^2^, sex, fluid intelligence, pF, pF_2,_ motion and scanning site.

## Results

We uncovered the same pattern of dFC between networks as previously reported (Fig 2a, Schwab et al., 2018), when using only data from independent subjects that were not used in Schwab et al. (2018) (Fig 2b) and likewise when using all available HCP data (Fig 2c). The cerebellar and auditory network appeared to be mostly a receiver in terms of directional information flow in the network.

### Significant effects of sex and motion on directed connectivity

Analysis of edge-wise associations of dFC with age, age^2^, sex, intellectual abilities, mental health and motion in the full HCP sample (N=984) yielded significant effects after Bonferroni correction (Figure 3). The findings show that compared to women, the VM in men receives less information from other visual networks, VO and VL (fig.3; SI tables 1a-b provides z-scores and corresponding p-values). Furthermore, the DMN-VM and DMN-VL edges was found to be less present in males, and the same pattern was found for the Ex-SM edge. In addition, motion had significant impact on directed connectivity between the Cer and VM and for the FPL-Ex edge, whereas age, age^2^, intellectual abilities and mental health, were not significantly associated with directed connectivity at the edge-level.

### Node-level analysis reveals significant effects of age^2^, sex and motion on directed connectivity

Next, we assessed out-degree and in-degree for networks. In line with results from the edge-wise analyses, we found that sex was significantly associated with out-degree (fig.4a), with networks in general sending less information in males compared to females. In addition, there was a significant relationship between the VM node and age^2^, where we observed a higher out-degree with higher age, indicating that the VM node sends more information as an effect of aging. Moreover, when looking at the in-degree (fig.4b) we also found significant effects of sex. Specifically, the VM, SM and Ex sends less information to other nodes in males compared to females, the same pattern as shown for sex effects and out-degree. In addition, motion was also associated with out-and in-degree for the Cer and Ex nodes. Further, we did not find an effect of age, cognition or mental health on node-level.

### Similar investigations in older individuals revealed effects of age, sex, motion and scanner on dFC

Next, we employed the same analysis approach using UK Biobank data (age range: 45-80 years). We partly found similar patterns of dFC between networks as previously reported by Schwab et al. (2018). Whereas the characteristic of the Au network to have many input connections as found in HCP data did not replicate, UK Biobank data confirmed this pattern for the cerebellum, as well as a bidirectionality of the VM-VL edge with these nodes having a reciprocal information flow (Fig. 5).

Edge-wise analysis of dFC alterations related to age, sex, cognition, psychopathology, motion and scanning site is illustrated in Figure 5 (SI tables 2a-e provides z-scores and corresponding p-values for UK Biobank data).

We found a significant effect of age on edge-wise information flow with a positive association for the VL, and cerebellar network, with these nodes giving more information to the DMN with higher age. Further, with higher age the DMN gives more information to the VL and Cer network, while the Ex receives more information from the FPR, FPL and the SM (Fig.5; see SI for further details). Moreover, SM receives less information in general from the other nodes and this node sends less information input in the information flow with the cerebellar and auditory networks with higher age. In addition, VM sends less information to the cerebellar network and there was also a decrease in information flow from VO to FPL, VL-FPL and FPR-FPL, FPL-FPR, and for the DMN and VL to the Au network. In addition, there was an effect of age^2^, from Cer to the FPR node.

Also, there was widespread significant associations between dFC and sex (fig.5; see SI for further details), where the FPR, FPL, SM, Cer, Au and Ex nodes in males more often receive information in general from the other nodes compared to females. The opposite was found for VO-VM, and there was bidirectional dFC between DMN and Cer with reduced information flow in both directions observed in males. The opposite reciprocal relationship was found between the DMN and VM, with increased information flow present in males. Also, the DMN in males received more often information input from Au and VO compared to females, while the SM node in males more often sent information to the VO. In addition, the VL received more information from the VO, SM and Cer in males. The supplementary information covers results from additional analyses of interaction effects (SFig.3-4) and the impact of scan duration between HCP and the UK Biobank sample (SFig. 1-2).

### Node-level analysis reveals significant effects of age, age^2^, sex, mental health, motion and scanner in directed connectivity

When looking at the out-degree, or the number of output connections for a given node and the association with pF, we found that the visual networks as well as the SM and Au showed a negative association, with a higher number of output connections being related to a lower degree of depressive/anxiety symptoms (fig.6a). In addition, males showed a stronger pattern of nodes sending more information in general to the other networks compared to females. Also, in relation to age, the VO and VM had a negative association with out-degree, while age^2^ in general had a negative association with less output connections with higher age. Further, when estimating in-degree, or the number of input connections for a given node, males showed more marked receiver nodes than females for all the nodes, with the exception of the VO and VM that did not show an effect of sex (fig.6b). There were likewise effects of both age and age^2^, where specifically the FPR showed a positive association with age^2^, while the same was found for age and Ex in addition to opposite effects that were observed for the FPL, SM and Au and age. Motion was found in general for all nodes in both node-level analysis, and scanner also had a significant relationship with nodes in general except for Au and VO (for assessment of the balance between in-degree and out-degree see SI, SFig. 5-6).

**Figure 6:**
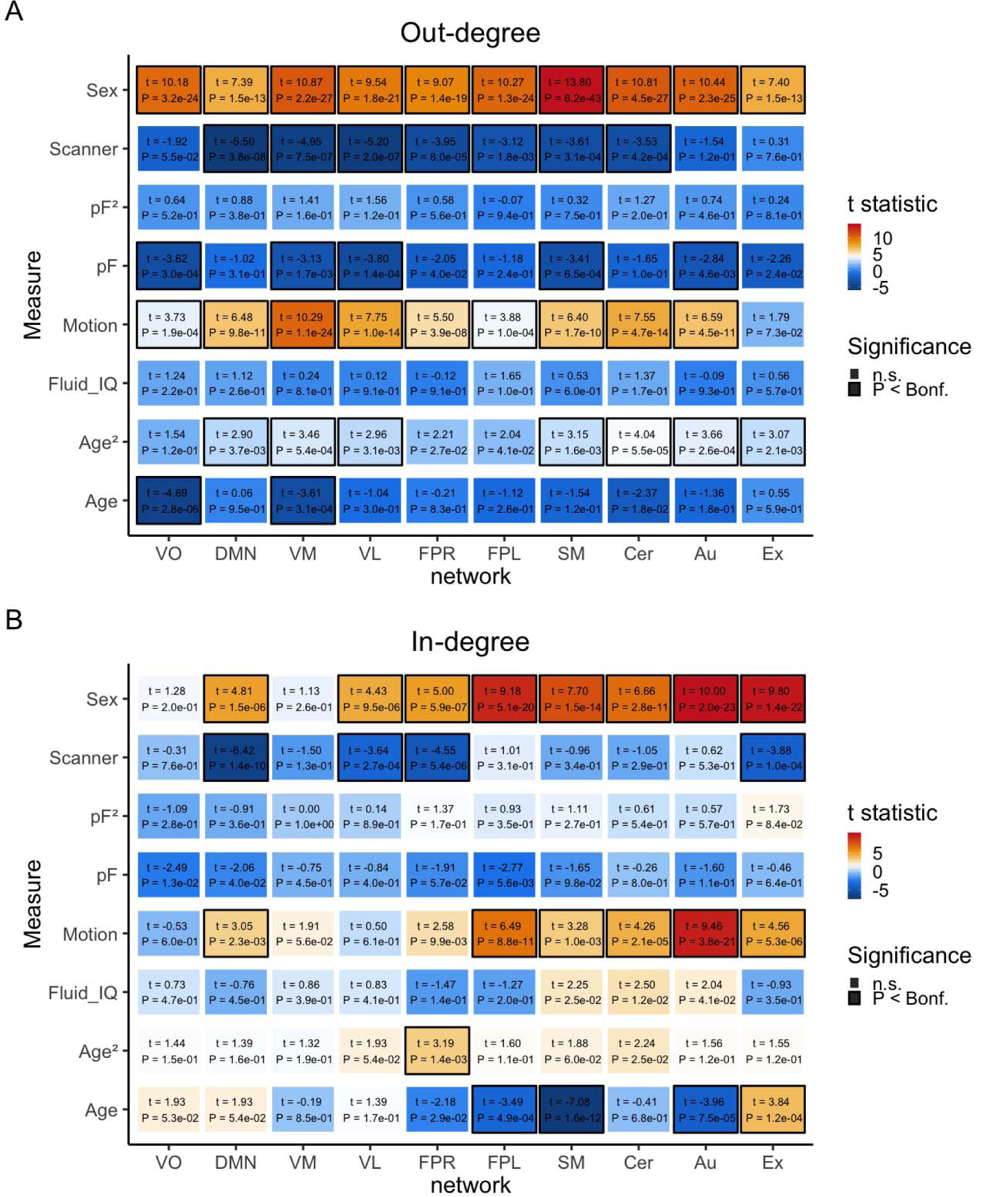
A) Out-degree matrix with corresponding effects of covariates age, age^2^, sex, fluid intelligence, pF, pF_2_, motion and scanner for UK Biobank (N=10,249, 45-80 years, df=10240). B) In-degree matrix with corresponding effects for the same covariates as in panel A. The colors reflect the t-value for the corresponding effect where red indicates a positive association and blue a negative association. Numbers inside the boxes indicate t-statistic and p-value, where significant effects are marked with a black border following Bonferroni correction (p<0.05).

## Discussion

The aim of the current study was to test for associations between dFC and age, age^2^, sex, cognitive abilities and mental health between core brain networks after testing the reproducibility of the DGM approach. We performed the analysis in healthy participants from two large public cohorts that differed in their age range (HCP: 22-37 years, n= 984, UK Biobank: 45-80 years, n=10,249) including subjects displaying subclinical symptoms in psychiatric domains. Accordingly, by utilizing these healthy samples in our analysis, we are able to examine mental health on a continuum rather than treating it as a static condition in the population.

Using the same HCP subjects as Schwab et al. (2018) initially reported on, as well as independent HCP data, we replicated the patterns of dFC between networks. As seen in figure 2a, there were some minor differences from the patterns reported by Schwab et al. (2018) that can be attributed to the difference in ICA decompositions, as we utilized data processed in a newer HCP pipeline release than Schwab et al. (2018), where a different ICA decomposition was included. However, the similar patterns with different decomposition illustrate the robustness of the method. In addition, we investigated dFC in UK Biobank data. Both the HCP and UK Biobank samples confirmed that the cerebellar network receives mostly rather than emits information from several other networks. Further, the visual areas VM and VL showed a bi-directionality in the information flow of their connectivity, with effects particularly pronounced in UK Biobank. Whereas the previously reported patterns by Schwab et al. (2018) where the auditory network mostly receives information from the other network nodes replicated in the independent HCP analyses in the present study, similar patterns were not observed in UK Biobank data. These differences as well as the more symmetric findings in UK Biobank data may be attributable to sample specific distinctions, such as the differences in the age range or for instance dissimilarities in the decomposition of networks, or differences between the preprocessing pipelines used in the two samples.

We observed marked effects of age on dFC in UK Biobank sample. For example, the sensorimotor network generally received little information from other networks with higher age in the 45-80 years age range. This sensorimotor association with age is particularly interesting given that apparent aging of the brain appears a key characteristic in several common brain disorders (Hajek et al., 2019; Kaufmann et al., 2019; Schnack et al., 2016), including schizophrenia, which has repeatedly been associated with dysconnectivity of sensorimotor networks (Cheng et al., 2015; Kaufmann et al., 2015). Thus, it will be of interest to delineate age trajectories of dFC in mental disorders in future studies.

Moreover, the age effects were overall in the direction of decreased reception with higher age. However, three connections showed a bi-directional relationship with age with decreased connectivity flow in both directions between these nodes (Cer-SM, Au-SM and FPL-FPR). Additionally, two connections of the DMN increased bi-directionally with age (Cer-DMN, DMN-VL). Of note, increased connectivity between the cerebellum and the DMN with age has previously been reported in a study comparing a group of young to a group of old individuals using a static connectivity approach (Dørum et al., 2017). While connectivity was lower in the young group during rest, it was higher in the young group during task load (Dørum et al., 2017), which is in line with the established decline of DMN variability in old age (Maglanoc, Kaufmann, Jonassen, et al., 2019; Mowinckel, Espeseth, & Westlye, 2012). While direct comparisons between results obtained with dFC and those with static connectivity warrant caution given that they measure different properties of the rsfMRI timeseries, these results may suggest that changes in direction with age may also depend on task load. This will need to be explored in future studies where task-fMRI could be used to constrain dFC to specific states and task performance.

There were significant sex differences at the edge- and node-level, where for instance males showed a stronger pattern of nodes sending and receiving more information in general to the other networks compared to females in the UK Biobank sample. There was also a pronounced effect of sex on dFC in the sensorimotor network in UK Biobank data, with males showing a more marked pattern of dFC compared to females on the edge-level. Prior research using static functional connectivity estimates have reported increased connectivity in males in the sensorimotor network in resting-state (Scheinost et al., 2015) and both increased and decreased down regulation between males and females while participants were performing a motor task (Lissek et al., 2007). There were also significant effects of pF in relation to visual networks as well as the SM and Au, where a higher number of output connections was related to a lower degree of depressive/anxiety symptoms. As such, our findings can complement static functional connectivity estimates and be of help in yielding insight into how sex, age and psychiatric symptoms factor into information flow of large-scale brain networks. Additionally, this can give a better understanding of the connectome in general, and also in relation to sex differences found in symptom onset and burden in mental disorders.

In regards to the thresholding used in this method, the sparsity is to some degree an artificially induced term and while it can reflect that there is indeed no connection between two nodes, it can also mean that there is no strong enough connection between two nodes. Conceptually this makes a big difference from an anatomical standpoint. However, if we think about it as a way of thresholding the data to a degree that reveals strong patterns but dampens the noisier, less clear patterns, it conceptually reminds of the procedure in regular static functional connectivity analysis where regularization is deployed to dampen low correlations and to pronounce strong ones. Another aspect to consider, is that the relatively limited number of nodes included in this analysis may have hampered the detection of sending nodes and as such future analyses deploying more fine-grained network parcellations is warranted.

Whereas our results revealed distinct effects of age, age^2^ and sex on dFC on edge-level, and age, age^2^, sex and pF on the node-level, none of our analyses identified significant relations with individual differences in cognitive test performance, and we did not find a significant association of mental health on dFC in the HCP sample or for pF^2^ in the UK Biobank sample. Of note, we here studied variations in mental health in *healthy* individuals, where effects may be subtle compared to studies including patient data. For example, studies looking at differences between healthy individuals and patients with psychiatric disorders have observed alterations in connectivity direction (Lu et al., 2012; Rolls et al., 2018; Schlösser et al., 2003; Shannon et al., 2009; Wicker et al., 2008). Another factor that may have contributed to the lack of associations with mental health in the current study may be inherent in the tools taken to assess mental health. The MHQ in UK Biobank was taken a long time after the scanning and it may thus not be a solid marker of the state at the participants’ time of scanning. Likewise, due to differences in available data, we used different approaches for measuring mental health, estimating two principal components in UK Biobank and utilizing a sum score in the HCP data. The NIH Toolbox Cognition Total Composite Score used in HCP cuts across various cognitive domains and as such may not be sensitive to specific higher-order networks, and could indicate why we did not find an association between intellectual abilities and information flow between networks. However, studies have shown that there is an association with subdomains of this test and effective connectivity in nodes such as the frontoparietal network (Harding, Yücel, Harrison, Pantelis, & Breakspear, 2015). Also, the ASR item used to measure psychiatric and life function in HCP may not be specific enough as it represents a sum score of a range of domains extending to depression and anxiety, aggressive behavior, attentional problems and hyperactivity, personality traits, psychotic and abnormal behavior, risk taking and impulsivity, somatic complaints, and substance use.

## Limitations

The current study does not come without limitations. The data was processed in different pipelines and we thus chose not to analyze the two samples together as would have been of interest for studying age effects across the lifespan. While we observed various patterns across the two independent cohorts, there were also marked differences that might be partly attributable to confound effects, such as variability in the ICA decompositions, scanning site and motion.

A major challenge in estimating directed functional connectivity has been the influence of regional differences in hemodynamic lags (Chang, Thomason, & Glover, 2008; Wei, Liao, Yan, He, & Xia, 2017). Granger Causality, a time domain approach which was widely used before for estimating directed functional connectivity, has been criticized for being highly influenced by such lags (Smith et al., 2011). DGM reflects instantaneous relationships and does not solely consider lagged relationships like Granger Causality does. To find out how DGM is influenced by differing hemodynamic lags in various brain regions, Schwab et al. (2018) examined how lags in the hemodynamic response could potentially influence the DGM estimation of directed functional connectivity by simulating systematic lags of the hemodynamic response and found DGM performed well in these network simulations. It was observed that large lags that are still physiologically plausible do not introduce spurious relationships with DGM as may be expected with Granger causality, however, sensitivity of DGM can drop to detect such relationships.

Moreover, DGM estimates binary connections, which may have rendered the association analyses less sensitive. In addition, DGM requires high-quality fMRI data with a low TR and benefits from a high number of observations. The long scan duration needed to acquire such data may have increased the chance that participants may fall asleep while they are being scanned. This is especially a challenge for the HCP project were participants are in the MRI scanner for a long time period (Glasser et al., 2018; Liu et al., 2018). Finally, as noted above, the lack of strong variations in mental health in these healthy samples may have limited the ability to identify associations with mental health measures. Future research, involving patients with psychiatric disorders may reveal if and how information flow is associated with disorders or related to specific symptoms.

## Conclusions

In conclusion, using independent rsfMRI data from HCP as well as the UK Biobank samples we replicated several of the directed connectivity patterns from the original HCP analysis (Schwab et al., 2018). In particular, we observed a marked characteristic of the cerebellar network to receive information from many other networks, and found a bi-directionality in the information flow between the visual areas VM and VL. Further, there was widespread age and sex effects on information flow, where strong age effects where observed in the sensorimotor network. In addition, we found associations of mental health on information flow for the sensorimotor network as well as the visual and auditory nodes. Our findings support the use of DGM as a measure of directed connectivity in rsfMRI data and uncovered new insight into the shaping of the connectome in aging. Future studies should examine dFC in other samples and look at directional changes in connectivity in relation to clinical populations and in broader age ranges.

## Data and code availability

The data incorporated in this work were gathered from the Human Connectome Project and the UK Biobank resources. Software needed to estimate directed connectivity is available at https://github.com/schw4b/DGM.

## Acknowledgements

We thank Tom Nichols for advice and input on this work. This research has been conducted using the UK Biobank Resource (access code 27412, https://www.ukbiobank.ac.uk/) and using data provided by the Human Connectome Project, WU-Minn Consortium (Principal Investigators: David Van Essen and Kamil Ugurbil; 1U54MH091657) funded by the 16 NIH Institutes and Centers that support the NIH Blueprint for Neuroscience Research; and by the McDonnell Center for Systems Neuroscience at Washington University. This work was performed on the TSD (Tjeneste for Sensitive Data) facilities, owned by the University of Oslo, operated and developed by the TSD service group at the University of Oslo, IT-Department (USIT).

