## Supplementary Information for "Differences in directed functional brain connectivity related to age, sex and mental health"

**Content**

*Page*

1. Supplementary tables	1

2. Analysis of the impact of scan duration	5

3. Analysis of interaction effects on edge-level	7

4. Analysis of node balance	9

5. Analysis of interaction effects on node-level	12

**1. Supplementary tables**

|  | **VO** | **DMN** | **VM** | **VL** | **FPR** | **FPL** | **SM** | **Cer** | **Au** | **Ex** |
| --- | --- | --- | --- | --- | --- | --- | --- | --- | --- | --- |
| **VO** | NA | -0.32, p =1 | -3.73, p =0.01728 | -1.69, p =1 | -0.76, p =1 | -1.54, p =1 | -1.66, p =1 | 1.92, p =1 | -0.83, p =1 | -2.4, p =1 |
| **DMN** | -1.63, p =1 | NA | -3.62, p =0.02688 | -4.04, p =0.00491 | -0.53, p =1 | -1.85, p =1 | -2.25, p =1 | -0.42, p =1 | 0.16, p =1 | -2.62, p =0.78185 |
| **VM** | -0.92, p =1 | -1.12, p =1 | NA | -3.07, p =0.19165 | 0.18, p =1 | -0.92, p =1 | -1.63, p =1 | -0.14, p =1 | -0.62, p =1 | -2.32, p =1 |
| **VL** | -1.28, p =1 | -0.97, p =1 | -3.95, p =0.00709 | NA | 0.68, p =1 | -1.06, p =1 | -1.93, p =1 | -0.31, p =1 | -1.1, p =1 | -2.97, p =0.26825 |
| **FPR** | -0.87, p =1 | -0.58, p =1 | -1.69, p =1 | -0.63, p =1 | NA | -1.86, p =1 | -1.95, p =1 | -1.36, p =1 | -3.15, p =0.14735 | -2.03, p =1 |
| **FPL** | -1.05, p =1 | -1.2, p =1 | -2.89, p =0.3445 | -1.75, p =1 | -1.45, p =1 | NA | -1.8, p =1 | -1.25, p =1 | 0.16, p =1 | -2.21, p =1 |
| **SM** | -1.68, p =1 | -0.63, p =1 | -1.9, p =1 | 0.01, p =1 | -0.26, p =1 | -1.06, p =1 | NA | 0.47, p =1 | -1.84, p =1 | -2.81, p =0.45174 |
| **Cer** | -1.74, p =1 | -0.61, p =1 | -1.85, p =1 | -1.8, p =1 | -0.56, p =1 | -0.92, p =1 | -0.01, p =1 | NA | 0.35, p =1 | -1.58, p =1 |
| **Au** | -1.61, p =1 | -0.46, p =1 | -1.94, p =1 | -1.68, p =1 | 1.1, p =1 | -0.29, p =1 | -3.35, p =0.07291 | 1.95, p =1 | NA | -2.08, p =1 |
| **Ex** | -0.06, p =1 | -0.76, p =1 | -1.84, p =1 | -1.9, p =1 | 0.08, p =1 | -0.48, p =1 | -3.48, p =0.04473 | -0.6, p =1 | -2.33, p =1 | NA |

***STable 1a.*** HCP: Z and P_Bonf_ values for effects of sex on directed connectivity on the edge-level.

|  | **VO** | **DMN** | **VM** | **VL** | **FPR** | **FPL** | **SM** | **Cer** | **Au** | **Ex** |
| --- | --- | --- | --- | --- | --- | --- | --- | --- | --- | --- |
| **VO** | NA | -0.27, p =1 | 0.12, p =1 | 2.34, p =1 | 0.31, p =1 | 2.25, p =1 | -0.18, p =1 | 1.44, p =1 | -0.63, p =1 | 1.54, p =1 |
| **DMN** | -1.84, p =1 | NA | 0.99, p =1 | 1.54, p =1 | 0.54, p =1 | 0.69, p =1 | 0.33, p =1 | 1.62, p =1 | 1.07, p =1 | 1.9, p =1 |
| **VM** | -2.07, p =1 | -1.18, p =1 | NA | -0.38, p =1 | 0.56, p =1 | 1.12, p =1 | 0.89, p =1 | 3.82, p =0.0118 | -0.51, p =1 | 3.06, p =0.19724 |
| **VL** | -1.77, p =1 | -0.76, p =1 | -0.47, p =1 | NA | 1.13, p =1 | 1.5, p =1 | 0.17, p =1 | 3.22, p =0.11651 | -0.07, p =1 | 1.66, p =1 |
| **FPR** | -2.29, p =1 | -1.69, p =1 | -0.16, p =1 | 1.52, p =1 | NA | 0.41, p =1 | -0.88, p =1 | 1.26, p =1 | -1.41, p =1 | 1.56, p =1 |
| **FPL** | -0.85, p =1 | 0.05, p =1 | -0.54, p =1 | 1.45, p =1 | -0.43, p =1 | NA | 1.36, p =1 | 2.33, p =1 | -0.19, p =1 | 3.52, p =0.03957 |
| **SM** | -3.18, p =0.13318 | 0.38, p =1 | 1.73, p =1 | -0.19, p =1 | -0.11, p =1 | 0.4, p =1 | NA | 1.24, p =1 | -0.77, p =1 | 1.87, p =1 |
| **Cer** | -1.27, p =1 | -0.98, p =1 | 3.87, p =0.00968 | 2.41, p =1 | 0.16, p =1 | 1.51, p =1 | 1.29, p =1 | NA | 1.76, p =1 | 2.18, p =1 |
| **Au** | -2.31, p =1 | -0.26, p =1 | -0.01, p =1 | 2.17, p =1 | 1.59, p =1 | 0.85, p =1 | -1.32, p =1 | 3.07, p =0.18989 | NA | 3.33, p =0.07879 |
| **Ex** | -0.89, p =1 | -0.21, p =1 | -0.32, p =1 | 0.23, p =1 | -1.65, p =1 | 2.74, p =0.55415 | -0.19, p =1 | 0.94, p =1 | -1.17, p =1 | NA |

***STable 1b.*** HCP: Z and P_Bonf_ values for effects of motion on directed connectivity on the edge-level.

|  | **VO** | **DMN** | **VM** | **VL** | **FPR** | **FPL** | **SM** | **Cer** | **Au** | **Ex** |
| --- | --- | --- | --- | --- | --- | --- | --- | --- | --- | --- |
| **VO** | NA | -1.81, p =1 | -2.82, p =0.43227 | 1.14, p =1 | -2.79, p =0.48032 | -3.76, p =0.01516 | -6.24, p =0 | -0.97, p =1 | -1.52, p =1 | 0.91, p =1 |
| **DMN** | -0.65, p =1 | NA | 0.55, p =1 | 4.49, p =0.00064 | -0.31, p =1 | -3.45, p =0.05013 | -5.05, p =4e-05 | 5.21, p =2e-05 | -3.61, p =0.02802 | 2.74, p =0.55916 |
| **VM** | -0.67, p =1 | 0.24, p =1 | NA | -1.03, p =1 | -1.38, p =1 | -2.45, p =1 | -4.06, p =0.00439 | -4.72, p =0.00021 | -1.89, p =1 | 2.52, p =1 |
| **VL** | 2.33, p =1 | 4, p =0.00577 | -2.13, p =1 | NA | -0.74, p =1 | -3.84, p =0.01127 | -4.6, p =0.00039 | 1.5, p =1 | -3.56, p =0.03293 | 2.08, p =1 |
| **FPR** | 2.46, p =1 | 3.02, p =0.22379 | 1.28, p =1 | 0.02, p =1 | NA | -4.17, p =0.00269 | -3.09, p =0.17746 | -1.27, p =1 | -2.96, p =0.27717 | 4.08, p =0.00411 |
| **FPL** | 1.66, p =1 | 0.18, p =1 | 1.29, p =1 | -1.42, p =1 | -3.53, p =0.03781 | NA | -5.33, p =1e-05 | 0.42, p =1 | -1.66, p =1 | 3.94, p =0.00723 |
| **SM** | 0.57, p =1 | -0.88, p =1 | -0.78, p =1 | 2.26, p =1 | 1.58, p =1 | -1.58, p =1 | NA | -5.2, p =2e-05 | -5.77, p =0 | 3.63, p =0.02537 |
| **Cer** | 0.98, p =1 | 5.15, p =2e-05 | -3.07, p =0.19053 | 1.4, p =1 | -2.79, p =0.48123 | -2.18, p =1 | -9.01, p =0 | NA | -2, p =1 | 2.07, p =1 |
| **Au** | 3.2, p =0.1225 | -0.5, p =1 | 1.2, p =1 | -0.18, p =1 | -3.02, p =0.23019 | -1.61, p =1 | -7.75, p =0 | 2.39, p =1 | NA | 0.74, p =1 |
| **Ex** | 1.23, p =1 | 2.05, p =1 | 2.59, p =0.87327 | -0.76, p =1 | 0.45, p =1 | -0.36, p =1 | -3.32, p =0.07961 | 1.55, p =1 | -2.31, p =1 | NA |

***STable 2a.*** UK Biobank: Z and P_Bonf_ values for effects of age on directed connectivity on the edge-level.

|  | **VO** | **DMN** | **VM** | **VL** | **FPR** | **FPL** | **SM** | **Cer** | **Au** | **Ex** |
| --- | --- | --- | --- | --- | --- | --- | --- | --- | --- | --- |
| **VO** | NA | 0.16, p =1 | -0.53, p =1 | 0.45, p =1 | 1.6, p =1 | 0.2, p =1 | 0.67, p =1 | 0.45, p =1 | 1.7, p =1 | 0.71, p =1 |
| **DMN** | 0.76, p =1 | NA | 0.62, p =1 | 0.55, p =1 | 2.52, p =1 | 1.36, p =1 | 0.65, p =1 | -0.33, p =1 | 1, p =1 | 2.69, p =0.64405 |
| **VM** | 0.22, p =1 | 2.04, p =1 | NA | 2.07, p =1 | 2.29, p =1 | 1.02, p =1 | 0.19, p =1 | 2.75, p =0.53045 | 1.11, p =1 | 1, p =1 |
| **VL** | 0.79, p =1 | 0.74, p =1 | 0.73, p =1 | NA | 1.59, p =1 | 2.72, p =0.59214 | 1.46, p =1 | 1.31, p =1 | 0.48, p =1 | 0.72, p =1 |
| **FPR** | -0.91, p =1 | 1.61, p =1 | 1.24, p =1 | -0.03, p =1 | NA | 0.1, p =1 | 2.28, p =1 | 2.06, p =1 | 1.36, p =1 | 0.47, p =1 |
| **FPL** | 0.55, p =1 | 0.48, p =1 | 0.93, p =1 | 1.01, p =1 | -0.28, p =1 | NA | 2.32, p =1 | 0.11, p =1 | 1.27, p =1 | 0.73, p =1 |
| **SM** | 1.13, p =1 | 1.45, p =1 | -0.22, p =1 | 1.22, p =1 | 3.22, p =0.11637 | 0.22, p =1 | NA | 1.3, p =1 | 2.13, p =1 | 1.34, p =1 |
| **Cer** | 1.35, p =1 | 0.35, p =1 | 1.59, p =1 | 1.62, p =1 | 3.59, p =0.03024 | 2.17, p =1 | 1.73, p =1 | NA | 1.07, p =1 | 0.95, p =1 |
| **Au** | 2.79, p =0.47505 | 0.01, p =1 | 1.08, p =1 | 1.97, p =1 | 2.12, p =1 | 0.9, p =1 | 1.68, p =1 | 1.36, p =1 | NA | 0.6, p =1 |
| **Ex** | 1.61, p =1 | 1.53, p =1 | 0.53, p =1 | 0.89, p =1 | 1.82, p =1 | 2.21, p =1 | 0.92, p =1 | 1, p =1 | 0.02, p =1 | NA |

***STable 2b.*** UK Biobank: Z and P_Bonf_ values for effects of age^2^ on directed connectivity on the edge-level.

|  | **VO** | **DMN** | **VM** | **VL** | **FPR** | **FPL** | **SM** | **Cer** | **Au** | **Ex** |
| --- | --- | --- | --- | --- | --- | --- | --- | --- | --- | --- |
| **VO** | NA | 5.42, p =1e-05 | -3.89, p =0.00901 | 4.45, p =0.00078 | 4.28, p =0.00168 | 7.26, p =0 | 6.51, p =0 | 1.01, p =1 | 6.12, p =0 | 6.38, p =0 |
| **DMN** | 1.43, p =1 | NA | 6.17, p =0 | 0.06, p =1 | 1.27, p =1 | 5.72, p =0 | 4.15, p =0.00294 | -7.26, p =0 | 7.45, p =0 | 5.2, p =2e-05 |
| **VM** | -2.09, p =1 | 8.97, p =0 | NA | 0.17, p =1 | 5.69, p =0 | 7.54, p =0 | 4.94, p =7e-05 | 3.63, p =0.0258 | 7.04, p =0 | 2.66, p =0.69763 |
| **VL** | -0.19, p =1 | 1.77, p =1 | -1.59, p =1 | NA | 3.7, p =0.0197 | 5.4, p =1e-05 | 4.9, p =9e-05 | 7.64, p =0 | 3.64, p =0.02453 | 7.47, p =0 |
| **FPR** | 1.4, p =1 | 2.92, p =0.31951 | 1.28, p =1 | 2.17, p =1 | NA | 4.01, p =0.0054 | 6.25, p =0 | 3.65, p =0.02339 | 2.96, p =0.27656 | 7.19, p =0 |
| **FPL** | 1.48, p =1 | 2.63, p =0.76956 | 1.95, p =1 | -0.03, p =1 | 0.99, p =1 | NA | 5.81, p =0 | 8.05, p =0 | 6.79, p =0 | 8.42, p =0 |
| **SM** | 4.27, p =0.00175 | 3.16, p =0.14035 | 1.84, p =1 | 4.71, p =0.00022 | 6.58, p =0 | 8.4, p =0 | NA | 6.83, p =0 | 11.01, p =0 | 5.75, p =0 |
| **Cer** | -0.39, p =1 | -4.12, p =0.00339 | -0.21, p =1 | 7.65, p =0 | 4.57, p =0.00044 | 9.78, p =0 | 8.42, p =0 | NA | 9.39, p =0 | 5.18, p =2e-05 |
| **Au** | 1.33, p =1 | 5.84, p =0 | 1.65, p =1 | 0.07, p =1 | -2.43, p =1 | 7.29, p =0 | 8.59, p =0 | 5.61, p =0 | NA | 9.48, p =0 |
| **Ex** | 0.09, p =1 | 1.78, p =1 | -2.98, p =0.26036 | 1.59, p =1 | 4.06, p =0.00448 | 7.94, p =0 | 3.61, p =0.02788 | -0.49, p =1 | 10.11, p =0 | NA |

***STable 2c.*** UK Biobank: Z and P_Bonf_ values for effects of sex on directed connectivity on the edge-level.

|  | **VO** | **DMN** | **VM** | **VL** | **FPR** | **FPL** | **SM** | **Cer** | **Au** | **Ex** |
| --- | --- | --- | --- | --- | --- | --- | --- | --- | --- | --- |
| **VO** | NA | 0.96, p =1 | -1.78, p =1 | -1.57, p =1 | -0.07, p =1 | 3.94, p =0.0074 | 1.1, p =1 | 1.77, p =1 | 6.16, p =0 | 2.92, p =0.31249 |
| **DMN** | 0.25, p =1 | NA | 1.01, p =1 | 1.38, p =1 | 2.8, p =0.46279 | 4.56, p =0.00046 | 2.74, p =0.56022 | -0.89, p =1 | 8.07, p =0 | 1.78, p =1 |
| **VM** | -2.55, p =0.95585 | 4.25, p =0.00192 | NA | 0.35, p =1 | 4.78, p =0.00016 | 8.27, p =0 | 3.24, p =0.10615 | 4.9, p =9e-05 | 7.96, p =0 | 4.92, p =8e-05 |
| **VL** | -0.75, p =1 | 3.55, p =0.03488 | -1.93, p =1 | NA | 0.73, p =1 | 3.76, p =0.01512 | 3.77, p =0.01457 | 5.96, p =0 | 7.44, p =0 | 3.55, p =0.03445 |
| **FPR** | -1.36, p =1 | 2.13, p =1 | 2.1, p =1 | -0.28, p =1 | NA | 5.23, p =2e-05 | 2.11, p =1 | -1.93, p =1 | 6.99, p =0 | 3.46, p =0.04876 |
| **FPL** | -1.74, p =1 | 1.72, p =1 | 2.55, p =0.97298 | -3.21, p =0.12019 | 2.78, p =0.49305 | NA | 2.57, p =0.92671 | 2.09, p =1 | 6.32, p =0 | 0.57, p =1 |
| **SM** | -0.34, p =1 | 3.55, p =0.03428 | 1.82, p =1 | 1.03, p =1 | 0.71, p =1 | 5.08, p =3e-05 | NA | 4, p =0.00564 | 5.19, p =2e-05 | 3.41, p =0.05839 |
| **Cer** | 1.11, p =1 | -0.2, p =1 | 3.48, p =0.04585 | 4.84, p =0.00012 | -0.62, p =1 | 5.54, p =0 | 4.19, p =0.00249 | NA | 6.94, p =0 | 2.42, p =1 |
| **Au** | 2.55, p =0.96541 | 2.74, p =0.54887 | 1.27, p =1 | 0.3, p =1 | 3.41, p =0.05937 | 4.59, p =0.00041 | 1.9, p =1 | 2.66, p =0.71325 | NA | 3.86, p =0.01026 |
| **Ex** | -0.07, p =1 | -0.69, p =1 | -1.35, p =1 | -0.8, p =1 | 0.56, p =1 | 2.75, p =0.53621 | 0.97, p =1 | -0.34, p =1 | 5.1, p =3e-05 | NA |

***STable 2d.*** UK Biobank: Z and P_Bonf_ values for effects of motion on directed connectivity on the edge-level.

|  | **VO** | **DMN** | **VM** | **VL** | **FPR** | **FPL** | **SM** | **Cer** | **Au** | **Ex** |
| --- | --- | --- | --- | --- | --- | --- | --- | --- | --- | --- |
| **VO** | NA | -2.69, p =0.64304 | -0.14, p =1 | -2.48, p =1 | -0.27, p =1 | 0.5, p =1 | 0.13, p =1 | -0.97, p =1 | 0.16, p =1 | -0.99, p =1 |
| **DMN** | 0.14, p =1 | NA | -2.1, p =1 | -2.15, p =1 | -5.54, p =0 | 1.04, p =1 | -2.56, p =0.94996 | -3.71, p =0.01842 | 0.63, p =1 | -5.03, p =4e-05 |
| **VM** | -0.38, p =1 | -5.31, p =1e-05 | NA | 1.75, p =1 | -5.22, p =2e-05 | -0.82, p =1 | -1.69, p =1 | -1.74, p =1 | -0.78, p =1 | -2.25, p =1 |
| **VL** | -2.7, p =0.63349 | -3.15, p =0.14491 | 2.25, p =1 | NA | -2.72, p =0.59223 | -0.04, p =1 | -1.76, p =1 | -3.89, p =0.0091 | 0.68, p =1 | -5.68, p =0 |
| **FPR** | -0.37, p =1 | -6.28, p =0 | -2.04, p =1 | -1, p =1 | NA | 2.32, p =1 | -0.52, p =1 | -1.47, p =1 | 0.04, p =1 | -4.91, p =8e-05 |
| **FPL** | -0.51, p =1 | -3.53, p =0.03809 | -2.08, p =1 | -3.13, p =0.1557 | 0.39, p =1 | NA | -0.84, p =1 | 2.25, p =1 | 0.59, p =1 | -3.16, p =0.14238 |
| **SM** | 0.72, p =1 | -4.23, p =0.00213 | -2.07, p =1 | -3.09, p =0.18292 | -2.9, p =0.33081 | 0.22, p =1 | NA | -1.57, p =1 | 1.83, p =1 | -2.18, p =1 |
| **Cer** | -0.77, p =1 | -5.75, p =0 | -2.84, p =0.40281 | -4.17, p =0.00272 | -3.14, p =0.15355 | 1.92, p =1 | -1.05, p =1 | NA | 0.21, p =1 | 3.46, p =0.04899 |
| **Au** | 0.47, p =1 | -3.31, p =0.08446 | 1.49, p =1 | 0.78, p =1 | -2.98, p =0.25805 | 0.79, p =1 | 0.59, p =1 | -0.13, p =1 | NA | -2.63, p =0.77563 |
| **Ex** | 1.68, p =1 | -3.68, p =0.02104 | 1.92, p =1 | -3.19, p =0.126 | -4.2, p =0.00238 | 0.8, p =1 | 1.04, p =1 | 6.01, p =0 | 0.69, p =1 | NA |

***STable 2e.*** UK Biobank: Z and P_Bonf_ values for effects of scanner on directed connectivity on the edge-level.

**2. Analysis of the impact of scan duration**

To test if difference in paradigm acquisitions, namely if the length of the scan could influence the results between the two samples, we performed a supplemental analysis where we restricted the number of time points used in HCP to 490, as this was the identical number of observations as used in UK Biobank. We calculated the same associations as done for the main analysis in networks estimated from the restricted time series data. We found that reducing the number of time points had an effect on the connectivity patterns (figure S1). In consequence, the reported associations with sex for the VO-VM, DMN-VL and Ex-SM edge did not replicate (figure S2c). However, the VL-VM and DMN-VM connection which was associated with sex was replicated. There was also a significant association of dFC on the VM-Au edge for age^2^ (figure S2b). In addition, the Cer-VM edge was preserved but not the VM-Cer or FPL to Ex edge in relation to motion.

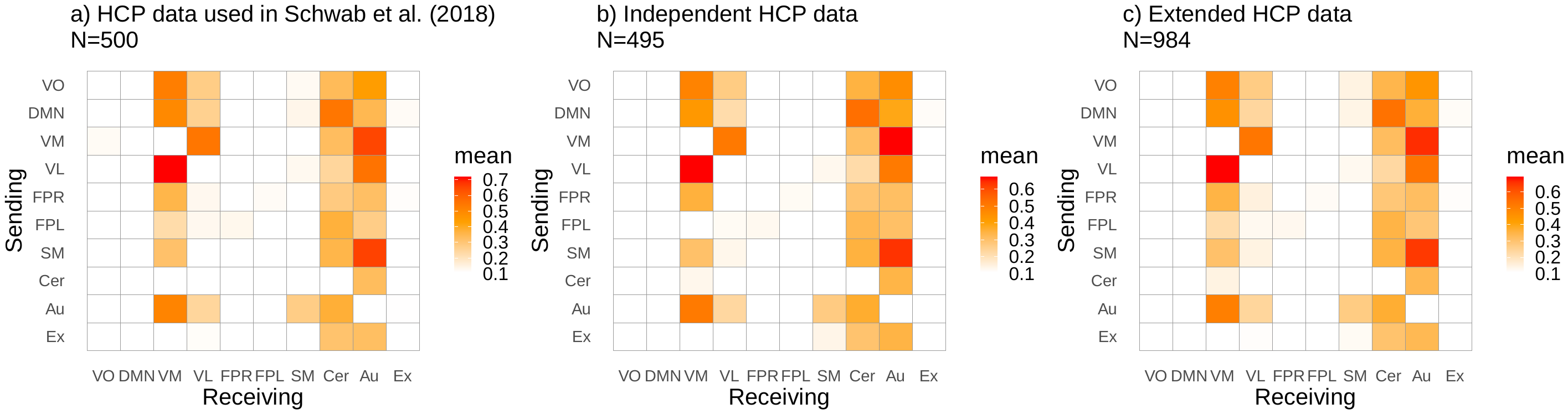

**Figure S1.** *Average directed connectivity matrices across subjects for HCP data showing the significant proportions of edges (binomial test, 5% FDR threshold) when including 490 time points in a) data previously reported by Schwab et al. (2018), b) independent data, c) all available data (a+b; slight differences in sample size due to differences in exclusion criteria). The legend shows the 10 RSNs included in the analysis; VO, visual occipital pole; DMN, default mode; VM, visual medial; VL, visual lateral; FPR, frontoparietal right; FPL, frontoparietal left; SM, sensorimotor; Cer, cerebellum; Au, auditory; Ex, executive control network, where the y-axis indicates the sender node, while the x-axis refers to the same nodes but here they are receivers.*

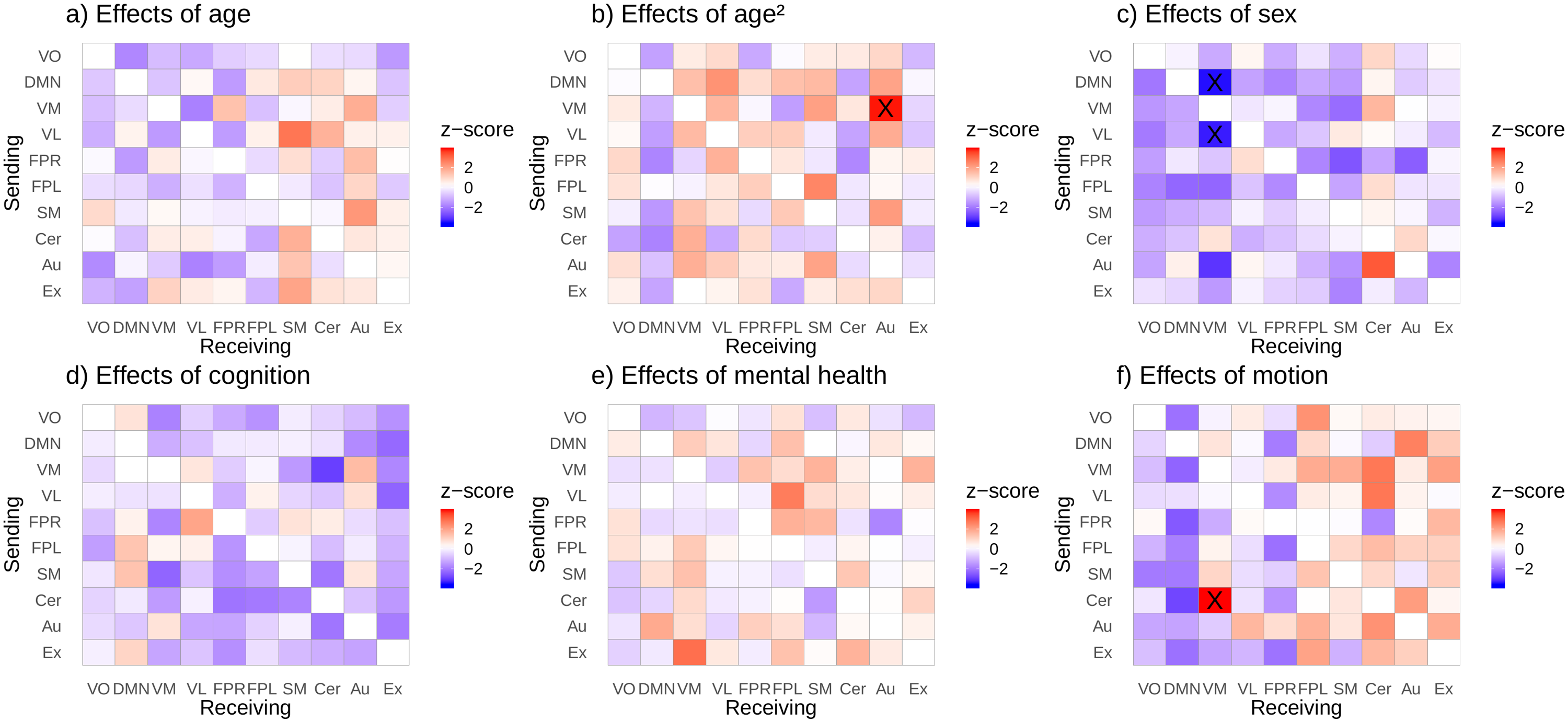

**Figure S2.** *Directed connectivity matrices showing the effects of age (a), age² (b), sex (c), intellectual abilities (d), mental health (e) and motion (f) on directed connectivity when including 490 time points. The analysis was performed in all available HCP data (N=984, 22-37 years). Significant edges following Bonferroni correction are marked as X. The y-axis indicates the sender node, while the x-axis refers to the receiving node. The colors reflect the z-value for the corresponding effects where red indicates a positive association and blue a negative association.*

**3. Analysis of interaction effects on edge-level**

In addition, we examined the interaction between sex and motion in both HCP (figure S3) and UK biobank data (figure S4). For the HCP sample, sex effects were no longer significant when adding an interaction term indicating a certain degree of motion dependence of the sex effects that were, however, not significant (none of the motion x sex effects with P<.05). For UK Biobank, including sex x motion in the model did have an impact on the number of sex effects, and there was one significant interaction effect of sex and motion on the Ex-VO edge, but overall there did not seem to be a relationship between sex and motion on directed functional connectivity. Given this and a non-significance of the interaction term for HCP, we found it appropriate to report statistics in the main manuscript from the model without the interaction term as it yielded a better fit.

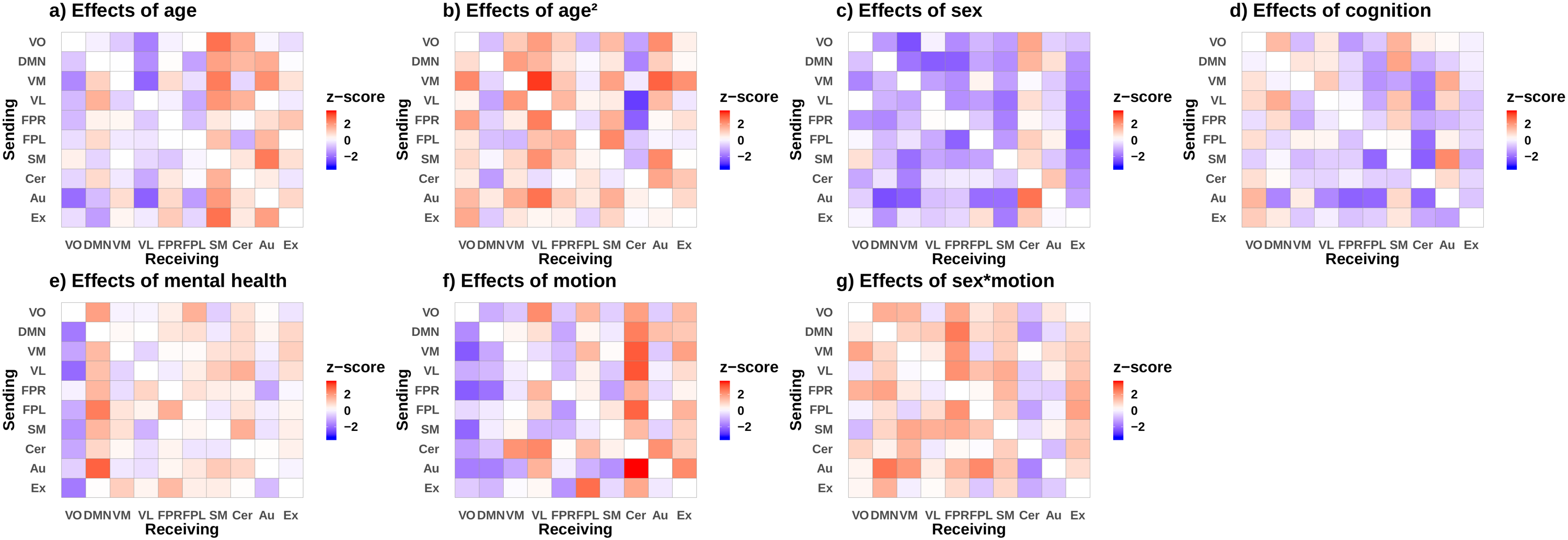

**Figure S3.** *Directed connectivity matrices showing the effects of age (a), age² (b), sex (c), cognitive abilities (d), mental health (e) motion (f), and sex x motion (g) on directed connectivity. The analysis was performed in all available HCP data (N=984, 22-37 years). Significant edges following Bonferroni correction are marked as X. The y-axis indicates the sender node, while the x-axis refers to the receiving node. The colors reflect the z-value for the corresponding effects where red indicates a positive association and blue a negative association.*

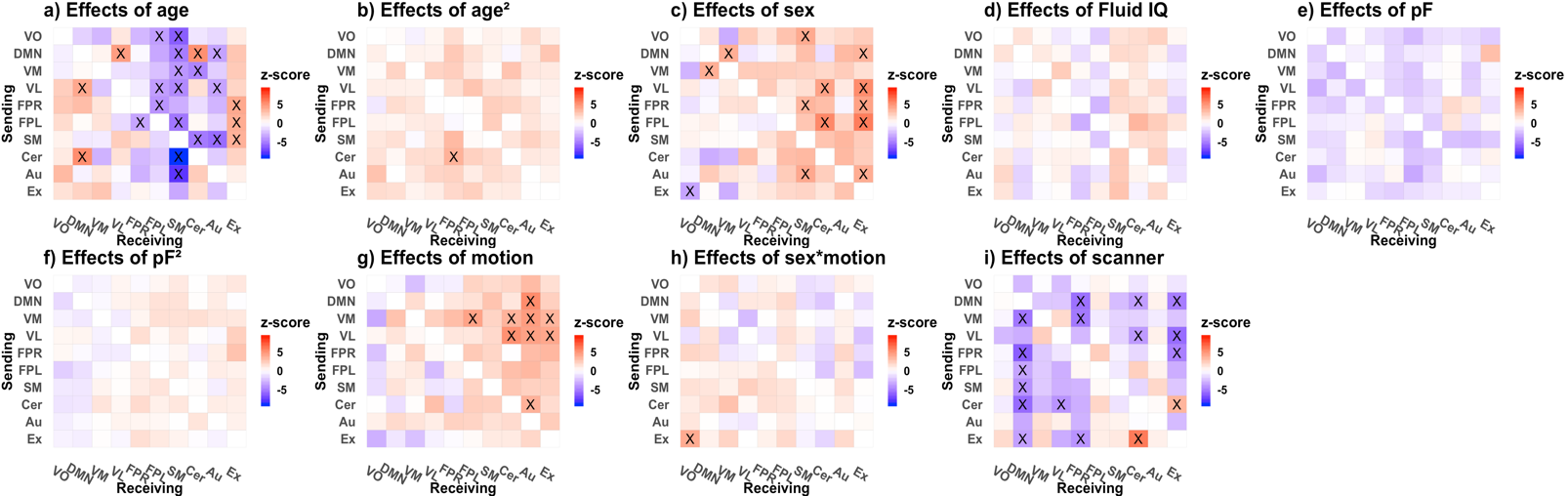

**Figure S4.** *Directed connectivity matrices showing the effects of age (a), age² (b), sex (c), fluid intelligence (d), pF (e), pF_2_ (f)_,_ motion (g), sex x motion (h) and scanner (i) for UK Biobank (N=10,249, 45-80 years). Significant edges following Bonferroni correction are marked as X. The y-axis indicates the sender node, while the x-axis refers to the receiving node. The colors reflect the z-value for the corresponding effects where red indicates a positive association and blue a negative association.*

**4. Analysis of node balance**

Furthermore, we assessed input and output connections for a given network together, to examine the balance between a network’s sent (out-degree) and received (in-degree) information. Accordingly, we calculated the ratio between the number of output connections and the number of input connections for a given node. To avoid inducing missing values when the denominator is 0, we added 0.5 to the denominator and nominator of dFC before taking the ratio (Sankey, Weissfeld, Fine, & Kapoor, 1996). We performed linear regression using this balance as a dependent variable and the same independent variables as used on the edge-level, where p-values for node balance were Bonferroni corrected for a number of 10 analysis, with an alpha level of 0.05.

Node-level analysis reveals significant effects of sex and motion on directed connectivity for HCP

Assessing network balance, we found that sex was significantly associated with node balance of the DMN and FPR network, with the balance in these networks decreased in males. In addition, there was a significant relationship between motion and the VO and DMN, where we observed a higher balance with higher motion. There were no significant effects of node balance for age, age^2^_,_ cognition or mental health.

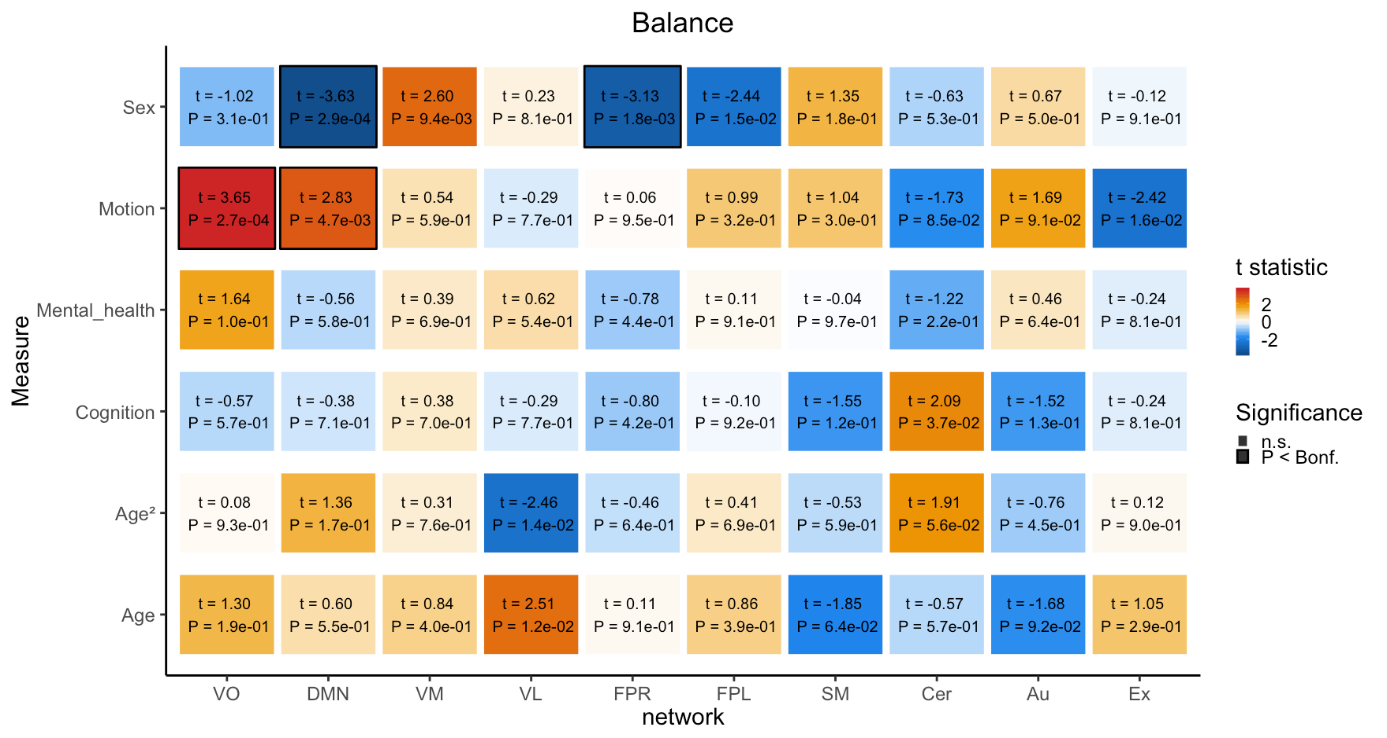

**Figure S5.** *Balance measure with corresponding effects of covariates age, age^2^, sex, cognitive abilities, mental health and motion for HCP data (N=984, 22-37 years). The colors reflect the t-value for the corresponding effect where red indicates a positive association and blue a negative association. Numbers inside the boxes indicate t-statistic and p-value, where significant effects are marked with a black border following Bonferroni correction (p<0.05).*

Node-level analysis reveals significant effects of age and sex in directed connectivity for UK Biobank

Further, we examined network balance in UK Biobank data (figure S6). Here we found several significant effects of age and sex. In brief, the VO and VM showed increased balance in males while the FPL, Au and Ex showed decreased balance in males. Further, the FPL, SM and Au showed increased balance with higher age, while VO and Ex showed decreased balance with higher age. In addition, we observed confound effects of scanning site on the DMN, FPR and Ex, and confound effects of motion on VM, VL, FPL, Au and Ex. We did not find any significant associations between node balance and age^2^, mental health or fluid intelligence. Taken together, these findings accompany the reported age and sex effects on in- and out-degree in the main manuscript, indicating that also the balance between in- and out-degree may be variant with age and sex.

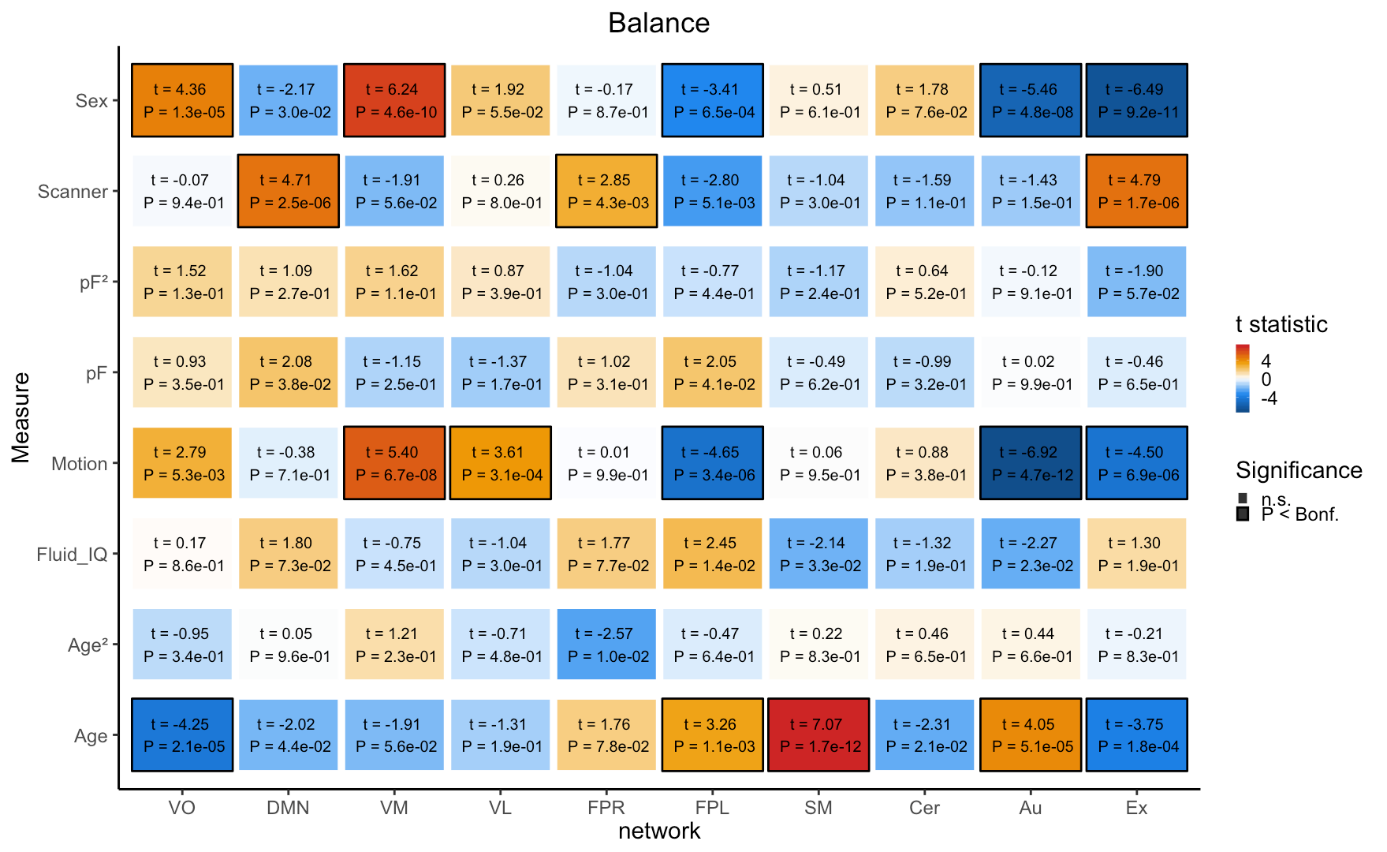

**Figure S6.** *Balance measure with corresponding effects of covariates age, age^2^, sex, fluid intelligence, pF, pF_2_, motion and scanner for UK Biobank (N=10,249, 45-80 years). The colors reflect the t-value for the corresponding effect where red indicates a positive association and blue a negative association. Numbers inside the boxes indicate t-statistic and p-value, where significant effects are marked with a black border following Bonferroni correction (p<0.05).*

**5. Analysis of interaction effects on node-level**

Analysis of balance and effects of interaction

Additionally, we also considered interaction effects of sex and age in relation to balance using the HCP (figure S7) and UK Biobank samples (figure S8). When including this interaction term in our model for each sample, the effects of sex on balance were no longer significant. This indicates that there is an influence of age on sex. However, we did not find a significant interaction between sex and age (alpha level = 0.05).

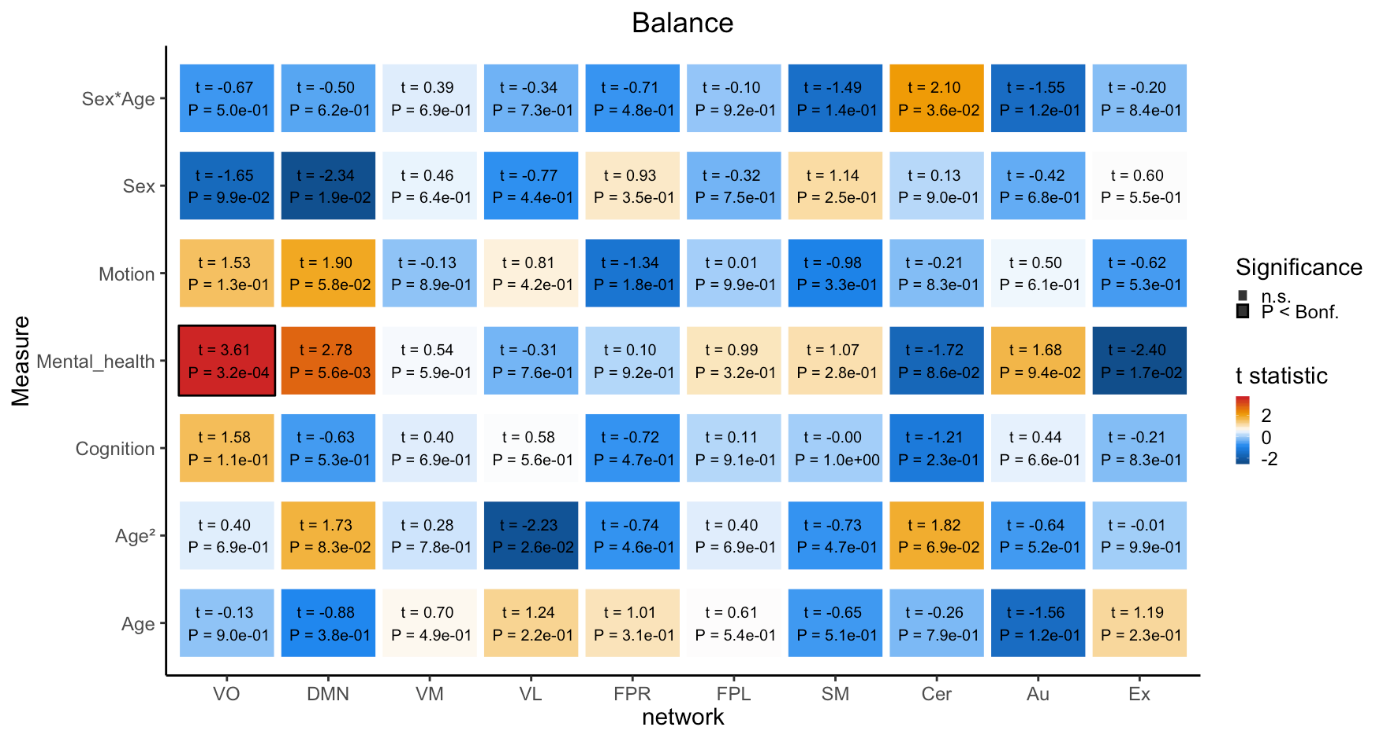

**Figure S7.** *Balance measure with corresponding effects of covariates age, age^2^, sex, sex x age, cognitive abilities, mental health, and motion for HCP data (N=984, 22-37 years). The colors reflect the t-value for the corresponding effect where red indicates a positive association and blue a negative association. Numbers inside the boxes indicate t-statistic and p-value, where significant effects are marked with a black border following Bonferroni correction (p<0.05).*

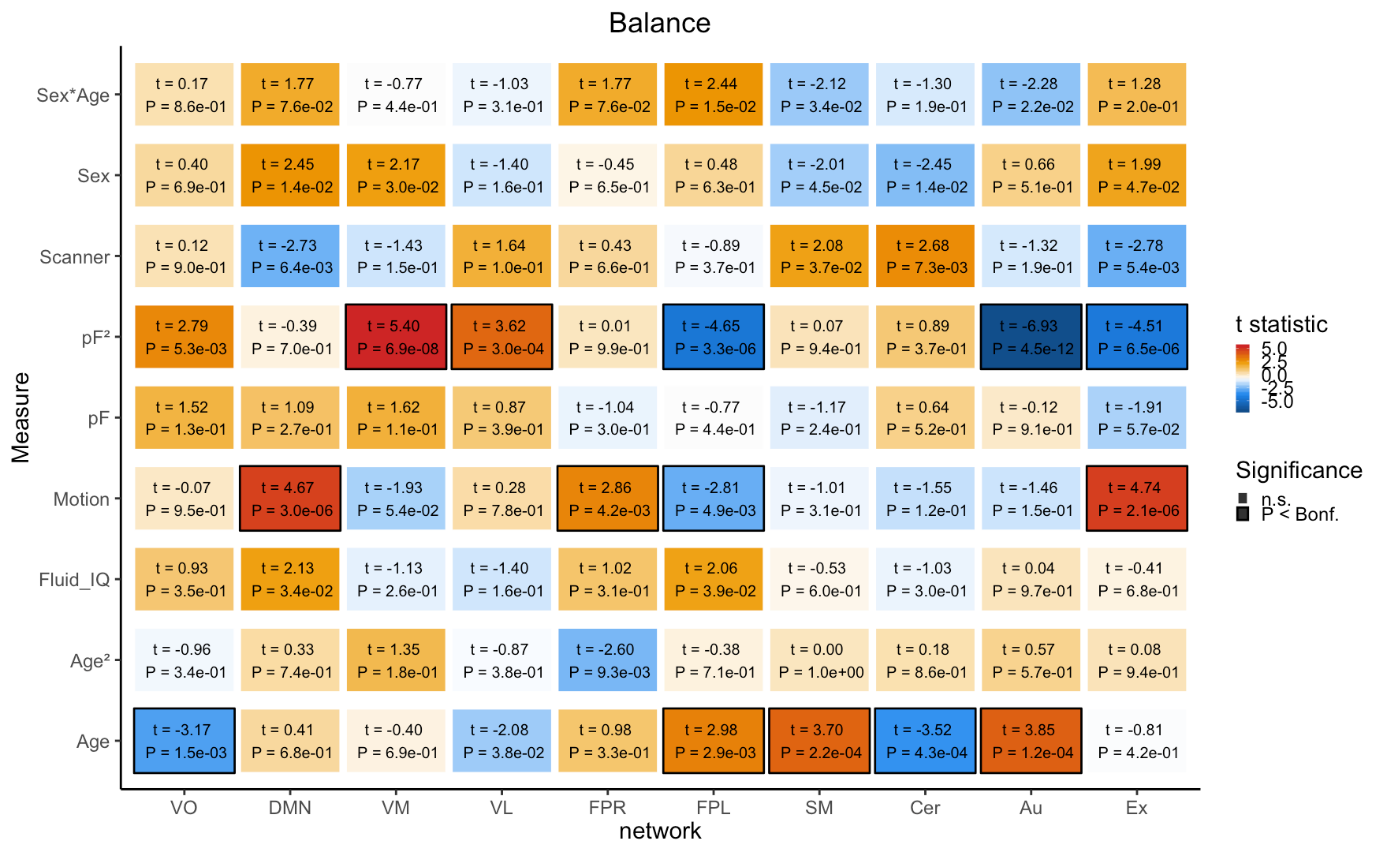

**Figure S8.** *Balance measure with corresponding effects of covariates age, age^2^, sex, sex x age, fluid intelligence, pF, pF_2,_ motion and scanner for UK Biobank (N=10,249, 45-80 years). The colors reflect the t-value for the corresponding effect where red indicates a positive association and blue a negative association. Numbers inside the boxes indicate t-statistic and p-value, where significant effects are marked with a black border following Bonferroni correction (p<0.05).*

Analysis of in-and out-degree and effects of interaction

Furthermore, we also considered interaction effects of sex and age in relation to in-and out-degree in both the HCP (figure S9) and UK Biobank samples (figure S10). For the HCP sample there were no significant interaction effect of sex and age on either out- or in-degree (alpha level = 0.05) and the effect of sex on both in-and out-degree were no longer significant when considering this interaction. Further, when including this interaction term in our model for UK Biobank, we found positive associations for sex and age on out-and in-degree for several nodes (age x sex effects with P<.05), and the effects of sex on out-and in-degree were no longer significant.

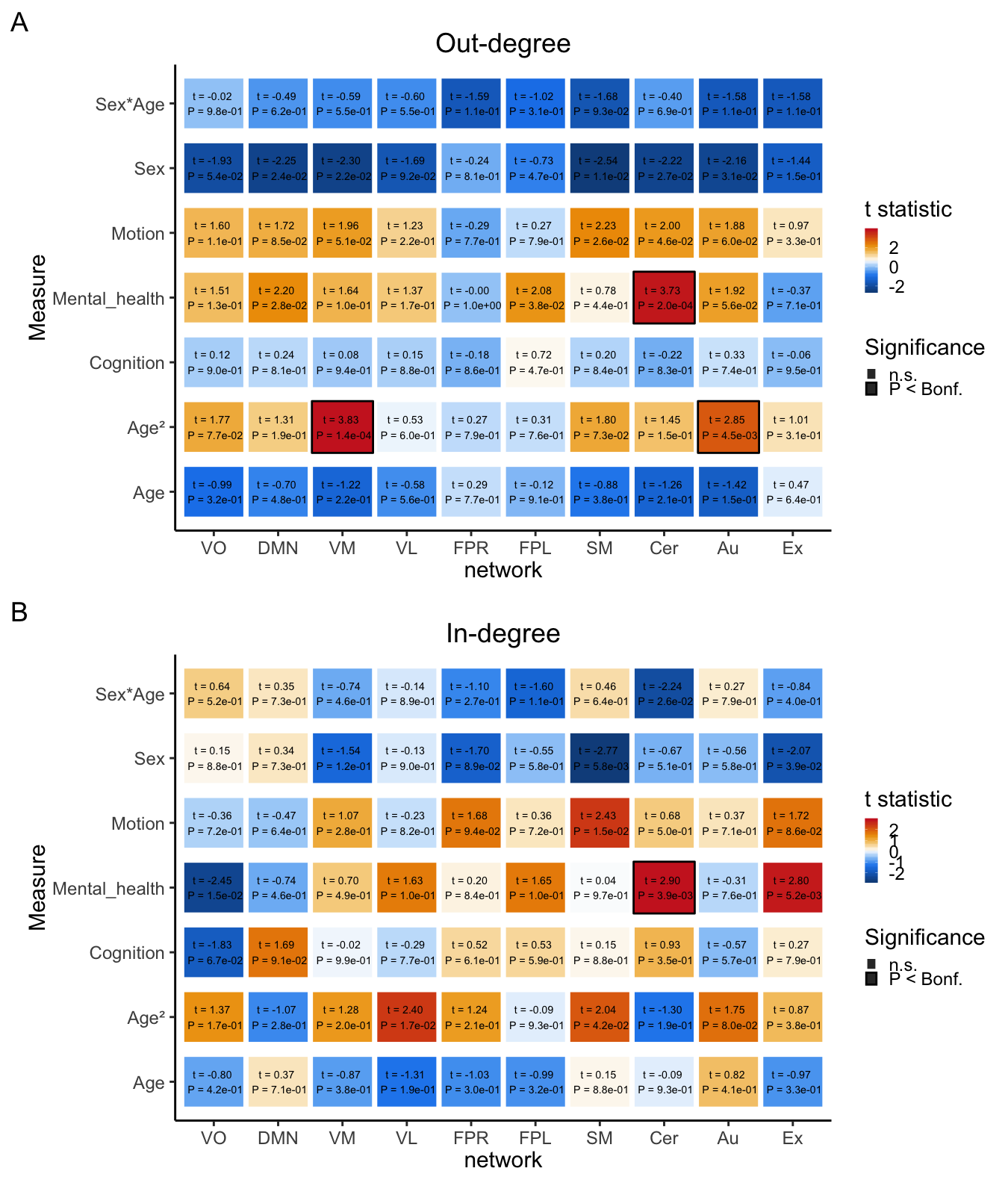

**Figure S9.** *Out-and In-degree measure with corresponding effects of covariates age, age^2^, sex, sex x age, cognitive abilities, mental health, and motion for HCP data (N=984, 22-37 years). The colors reflect the t-value for the corresponding effect where red indicates a positive association and blue a negative association. Numbers inside the boxes indicate t-statistic and p-value, where significant effects are marked with a black border following Bonferroni correction (p<0.05).*

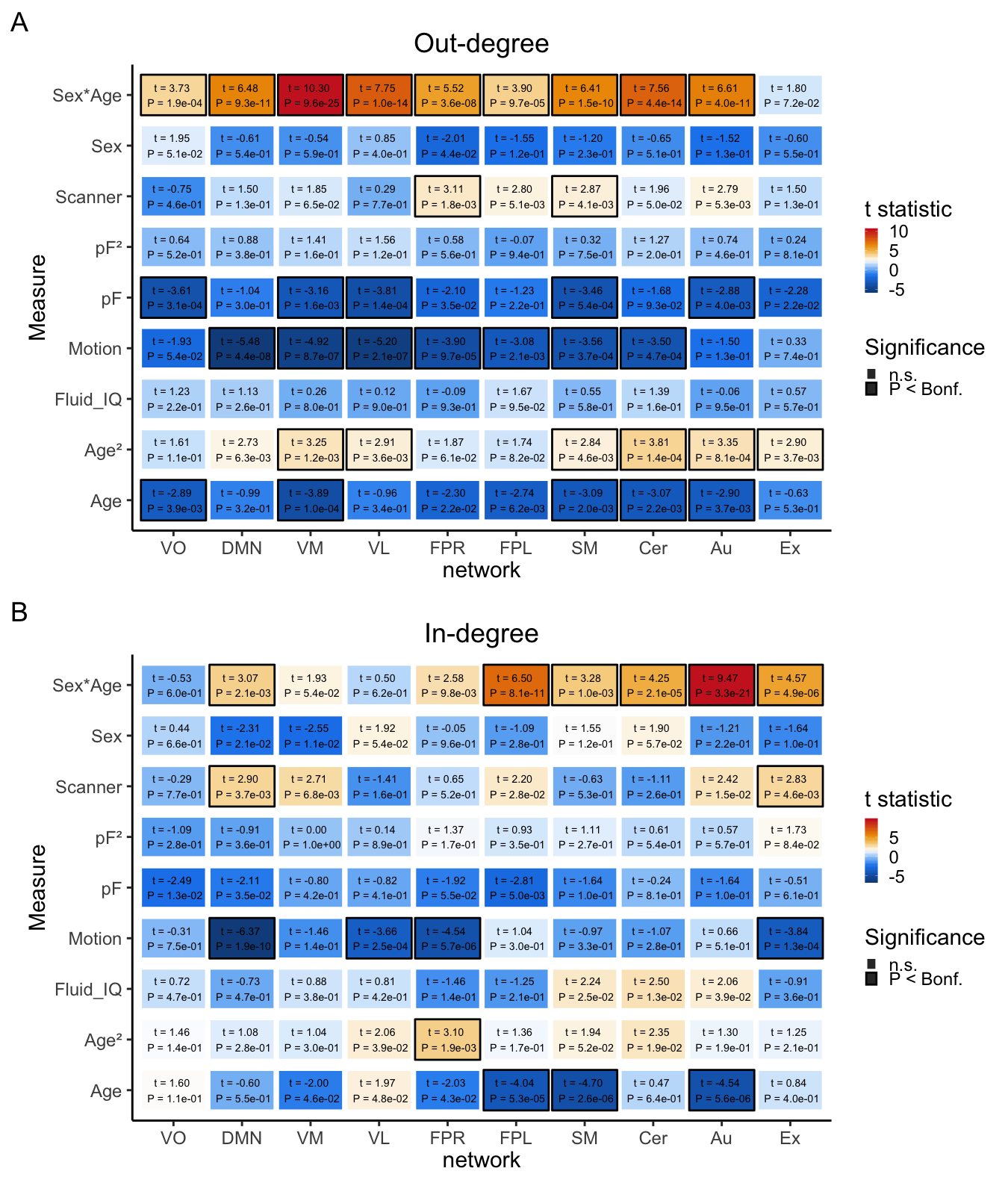

**Figure S10.** *Out-and In-degree measure with corresponding effects of covariates age, age^2^, sex, sex x age, fluid intelligence, pF, pF_2,_ motion and scanner for UK Biobank (N=10,249, 45-80 years). The colors reflect the t-value for the corresponding effect where red indicates a positive association and blue a negative association. Numbers inside the boxes indicate t-statistic and p-value, where significant effects are marked with a black border following Bonferroni correction (p<0.05).*
